# Multiple mutations acquired into canine RecQ-like helicases encoded by the aneuploid genome of transmissible sarcoma

**DOI:** 10.1101/508101

**Authors:** Wadim J. Kapulkin

## Abstract

Sticker sarcoma – a highly aneuploid, contagious neoplasm circulating in a domestic dog population - is broadly referred as a canine transmissible venereal tumour (CTVT). The karyotype of transmissible Sticker sarcoma appears as a collage of numerical and structural aberrations; the CTVT genome represents the generalized but stable neoplastic aneuploidy of monoclonal origins. Presented is an analysis of genetic events and variants underlying the aneuploid genomic structure of Sticker sarcoma described previously by Murchison et al. (2014) and Decker et al. (2015). Here we explored the above CTVT genomic compendia and mined the existing data - specifically looking for cases of convergence of multiple non-synonymous variants onto a single gene - the mutational patterns indicative for Knudsonian ‘two-hit’ kinetics. A Table I is given, providing theoretical estimates of retaining the intact wild-type copy, expected as a function of a cumulative mutational convergence observed in unphased sequence consensus. We demonstrate that the two canine RecQ-like helicases: Bloom syndrome helicase and RECQL4, encoded by the aneuploid transmissible tumour, have accumulated a multitude of different mutations. Among the sets of most intensely mutated transmissible sarcoma genes, we also identified a canine FANCD2 – yet another previously unnoticed multiple-hit candidate factor. We discuss a possible role of mutated RecQ-like helicases and other cooperating factors, perceivably involved in the maintenance of the neoplastic aneuploidy. We suggest the proposed cooperative actions of CTVT RecQ-like DNA helicases could be relevant interpreting whether variants contributing to RecQ-dependent karyotypic traits, respond to selective pressures that preserve the aneuploid genomic structure of transmissible Sticker sarcoma.

## INTRODUCTION

Cancer is a complex pathological process resulting in a common disease state occurring in animals including humans. This pathological process through which benign euploid cells acquire capabilities that cause its transformation into tumorigenic and eventually malignant cells. Typically, aberrant tumorigenic cells either activate eliminating mechanisms executing apoptosis and / or attract the stringency of the immune system. However, some transformed cells evade the immune response and programmed cell death mechanisms and will eventually establish and form clinically manifesting neoplasms. According to the multistage theory of tumorigenesis, transformed euploid cells eventually progress through uncontrolled growth towards malignant state resulting in an aggressive growth of a tumour (Weinberg R.A. 2014).

At the cellular level, the uncontrolled growth of initially euploid cells is often associated with changes to the karyotype broadly referred as aneuploidy. Aneuploidy is characterized by the imbalance to chromosome complement and is frequently attributed to the appearance of the aggressively proliferating malignant cells. Aneuploidy is regarded as clinically relevant in oncology as is often associated with unfavorable prognosis and can be an aid in distinguishing benign from malignant lesions. The hyper-proliferative properties of transformed cells leading to aneuploidy are observed in the affected tissues in a form of mitotic figures where some represent atypical mitoses (e.g. Therman & Timonen 1950; reviewed in Steinbeck 2001). Atypical mitoses (i.e. multipolar or other abnormal pathological mitotic forms) resulting in an uneven and altered distribution of the genetic material into daughter cells. Hyper-proliferating aneuploid cells subjected to constant and relentless replicative stress eventually progress into malignant forms. Neoplastic cells are genetically different from other (typically benign and stable) somatic cells of a host organism and progress into malignant states typically affected by some form of chromosomal instability. Oncogenic transformation associated changes leading to malignancies and aneuploidy, at sequence level, involve: gain- and loss-of-function alterations to individual genes; chromosome loss and gains; loss-of-heterozygosity (LOH) and somatic non-disjunction events; telomeric transactions including chromosome fusions, and other gross rearrangements. In particular aneuploid karyotype of neoplastic cells appear associated with loss-of-heterozygosity (LOH). At the base pair level, LOH is defined as a segmental allelic imbalance resulting from partial or complete somatic loss of one of the parental chromosomes. Somatic LOH is clinically relevant in oncology and is regarded as one of the mechanisms leading to complete loss-of-function of tumour suppressor genes (Weinberg R.A. 2014).

Malignant aneuploid cells acquire the ability to invade other tissues and organs (either through contact with locally aggressive tumour growth or various forms of metastases) of the host. The spread of aneuploid, malignant cells is typically restricted to the individual organism of an animal we refer to as a primary host and is limited by the primary hosts lifespan. Few neoplasms, however, have adapted an exceptional tropism to emerge from primary host organism to acquire a secondary host. Eruption and spawn of the contagious tumour cells from a primary host is assumed to proceed the eventual transmission between the other individuals of the same species. Those neoplasms are collectively referred as transmissible tumours (Eward & Kirpensteijn 2017), however, broadly referred as ‘natural transmissible tumours’ are rare. Presently among vertebrates there are three reported examples: Canine Transmissible Venereal Tumour (CTVT, a.k.a. Sticker sarcoma) (Novinsky 1876a, Novinsky 1876b, Novinsky 1877, reviewed by Shimkin 1955); Contagious Reticulum Cell Sarcoma of the Syrian hamsters (Ashbel R 1945, Brindley & Banfield 1961); and more recently Devil Facial Tumour Disease (DFTD, a.k.a. transmissible parasitic tumour of Tasmanian devil) (Loh R. et al. 2006, Hawkins et al. 2006, reviewed in Ostrander et al. 2016).

Given the established singular and exceptionally long natural history (Murgia et al. 2006; VonHoldt & Ostrander 2006), emergence of the tumour from genetically exploited host (reviewed in, Lindblad-Toh et al. 2005; Ostrander 2012; Dobson 2013; Davis & Ostrander 2014), the genetic structure of canine transmissible tumour have met the attention of others. Modern-day CTVT (Thorburn et al. 1968 and Sellyei et al. 1970) genome is known to occur in two main karyotypic forms: most common aneuploid form and rare form regarded as stable polyploid (presumably double-aneuploid). The karyotype of most common transmissible Sticker sarcoma form appears as a collage of numerical and structural aberrations, consistent with the appearance of pathological, atypical mitotic forms reported in cytological preparations (Krithiga et al. 2005, Stockmann et al. 2011). Collectively the CTVT genome is described as a state of generalized aneuploidy involving the derangement of all canine chromosomes. The aneuploid karyotype of present-day CTVT however, appears fairly consistent between a different tumour isolates - with chromosome complement of 57-59 (+/-2) and around 118 in stable polyploid suggestive for a the double-aneuploid complement. Compared with euploid canine (2n=78, with acrocentric autosomes) CTVT chromosome complement represents consistently aberrant karyotype of 13-17 metacentric and 42 acrocentric chromosomes (reviewed in Das & Das 2000 and Ganguly et al. 2013). Others (Murchison et al. 2014 and Decker et al. 2015) have extended the analysis of the haplotypic structure of the aneuploid Sticker sarcoma into base pair resolution thus providing the description of the genetic events the genome of CTVT have encountered. Specifically Murchison and colleagues, as an important step towards the Sticker sarcoma telomere-to-telomere assembly, compared the short-read sequencing data of two CTVT geographical isolates - modern-day aneuploid Sticker sarcoma samples from collected from two dogs: Australian and Brazilian. The elegant follow-up study, described by Decker and colleagues, have extended the above perspective and focused on the improved extraction and annotation of deleterious high-quality short-read sequence variants shared by both geographical tumour isolates. Referred later as the Decker set, the compendium of variations shared by both geographical isolates is likely representative of mutational patterns acquired into the prototype Sticker sarcoma circulating in the ancient canine population. The above variation compendium has extended and improved the base-pair resolution analysis of the genetic events the genome of CTVT has encountered. In particular, the shared pattern of seven hundred twenty-eight chromosome-to-chromosome translocations exerted on both tumours (Fig. 3B in Decker et al. 2015), were described as a part of the lexicon of 7338 structural rearrangements that underline the aneuploid karyotype of the prototype Sticker sarcoma. Decker set analysis of the aneuploid chromosomal complement of CTVT have also extended on the previous notion concerning the chromosome-wide prevalence of the loss-of-heterozygosity (LOH). Consistent with a previous notion (Murchison et al. 2014), a large proportion of the present day CTVT genome lacks the heterozygosity - Decker and collegues (2015) suggested the sequentiality to widespread LOH events encountered by transmissible Sticker sarcoma. Moreover, the elegant follow-up analysis confirmed the aneuploid CTVT have experienced heavy mutational loads during the exceptionally long natural history. In a comparative argument concerning the CTVT mutational loads, the follow-up analysis established the transmissible canine Sticker sarcoma genome have accumulated considerably more somatic mutations than most of the non-transmissible neoplasms reported in humans (Alexandrov et al. 2013).

The unprecedented catalogue of neoplastic variants encoded by the aneuploid transmissible canine tumour provided us with the distinct opportunity to explore the mutation accumulation patterns, acquired into a genome of the Sticker sarcoma. This carefully curated compendium of tumour genes includes variants responsible for adaptive traits, the neoplastic cells have perceivably required to survive in a niche provided by the host. Traits required by the neoplastic cells are of broad scientific and clinical interests, therefore justify the implementation of the model provided by aneuploid canine transmissible sarcoma into a field of comparative oncology. Here we explored the genome-wide prevalence of neoplastic variants indicative for cases of somatic composite heterozygosity represented among the Sticker sarcoma genes. The underlying hypothesis assumes that some of the prototype Sticker sarcoma genes have been altered by the sequential and independent genetic events, therefore cumulatively have acquired a multitude of different mutations.

## MATERIALS AND METHODS

### Source of the DNA sequencing data on the transmissible canine tumour samples

Data were derived from experimental sets of short-read sequences reported by Murchinson and colleagues (2014) were regarded as the original data source. Sets presented and by Murchinson and colleagues represent genome-wide paired-end Illumina HiSeq reads of two canine transmissible venereal tumours and their hosts collected in Australia and Brazil. Four experimental sets analyzed by Murchinson and colleagues were aligned with reference canine genome and consisted of two independently sampled transmissible aneuploid tumours: 24T and 79T; respectively collected from two geographically isolated diploid dogs – female host 24H and male host 79H^1^.

**Table I.** Defined as an absolute complement of the corresponding rates – the theoretical expectations - predict the second-hit candidates retain only 50% chance of attaining the out-of-phase *trans*-heterozygous Knudsonian inheritance model, while multiple-hit candidates retain out-of-phase likelihoods equal or greater than 75%. Conversely the chance of retaining of the intact wild-type allele - consistent with all-*cis*-heterozygous phase constellation - implicates the convergence of all variants onto a remaining allele, hence single haplotype variant phase.

| *N – mutation number* | *P-probability estimate*<br>*(exact value)* | *P-probability estimate*<br>*(rounded two decimals)* | *P%-approximate*<br>*percentage estimate* |
| --- | --- | --- | --- |
| N=2 | 0.5 | 0,5 | 50,00% |
| N=3 | 0.25 | 0,25 | 25,00% |
| N=4 | 0.125 | 0,13 | 12,50% |
| N=5 | 0.0625 | 0,06 | 6,25% |
| N=6 | 0.03125 | 0,03 | 3,13% |
| N=7 | 0.015625 | 0,02 | 1,56% |
| N=8 | 0.0078125 | 0,01 | 0,78% |
| N=9 | 0.00390625 | 0 | 0,39% |

**Table II.** Pearson correlation coefficient scores indicating strong (second-hit candidates) to moderate (multiple-hit candidates) positive correlation between the chromosome-wide distribution of multiply-mutated out-of-phase candidates and the number of autosomally encoded proteins (Decker at al. 2015 Table S2; and Method section for source values). Values correspond to data presentded as graphs in Fig.1 and 2.

| *Heteroallele candidate counts* | *value of R* | *P-Value*<br>*significant at $p < .01$* | *The value of $R^2$,*<br>*the coefficient of*<br>*determination* |
| --- | --- | --- | --- |
| Chr 1-38 autosomal hits (X excluded) | 0.8684 | < .00001 | 0.7541 |
| Chr 1-38 hits including X | 0.8055 | < .00001 | 0.6488 |
| Chr 1-38 autosomal second-hit (X excluded) | 0.8805 | < .00001 | 0.7753 |
| Chr 1-38 second-hit including X | 0.8257 | .000016 | 0.6818 |
| Chr 1-38 autosomal multiple-hit (X excluded) | 0.617 | .000037 | 0.3807 |
| Chr 1-38 multiple-hit including X | 0.5598 | .000211 | 0.3134 |

**Figure 1.**
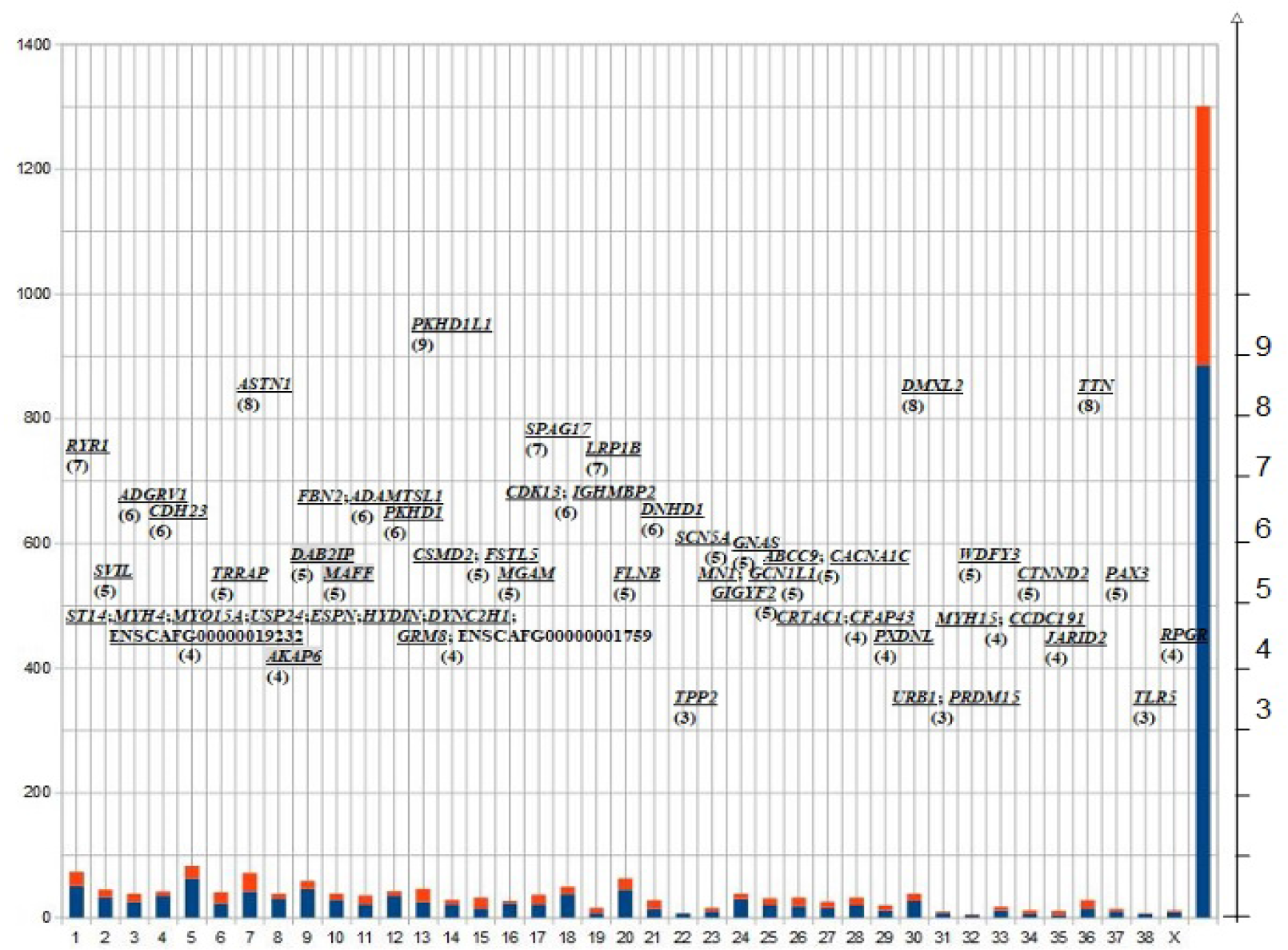
Genome-wide distribution of extreme examples among multiple-hit candidates. Canine chromosomes representation as vertical bar graph. Vertical bars correspond to each canine chromosome shown on x-axis (1-38 and X; last bar representing the cumulative value of total number of identfied candidates; areas corresponding to quantified second-hit and multiple-hit candidates are shown respectively in blue and red; quantified absolute values corresponding to number of identifed candidates representation by bar height show on y-axis to the left). Chromosome-wide distrbution of extreme examples of identified multiple-hit candidates. Individual CTVT loci are imposed onto the bar graph above the area corresponding to each canine linkage group. The number of hits recored for each corresponding extremally mutated candidate is indicated on y-axis drawn to the right but also indicated in brackets below the gene name (see Table II for details). Top ranking identification of CTVT-*PKHD1L1* (corresponding to referrence chr 13) accumulated nine mutations. CTVT-*SVIL* (chr 2), CTVT-GPR9S (chr 3), CTVT-AKAP (chr 8), CTVT-*DAB2IP* (chr 9), CTVT-FBN2 (chr 11), CTVT-*FSTL5* (chr 15), CTVT-*CSMD2* (chr 15), CTVT-SPAG17 (chr 17), CTVT-*CDK13* (chr 18), CTVT-*LRP1B* (chr 19), CTVT-GNAS (chr 24), CTVT-*GCN1L1* (chr 26), CTVT-*ABCC9* (chr 27), CTVT-CACNA1C (chr 27), CTVT-*CRTAC1* (chr 28), CTVT-*PXDNL* (chr 29), CTVT-*DMXL2* (chr 30), CTVT-*MYH15* (chr 33), CTVT-*CTNND2* (chr 34), CTVT-*JARID2* (chr 35) accumulated at least one additional structural variants (see Decker set SV for details).

**Figure 2.**
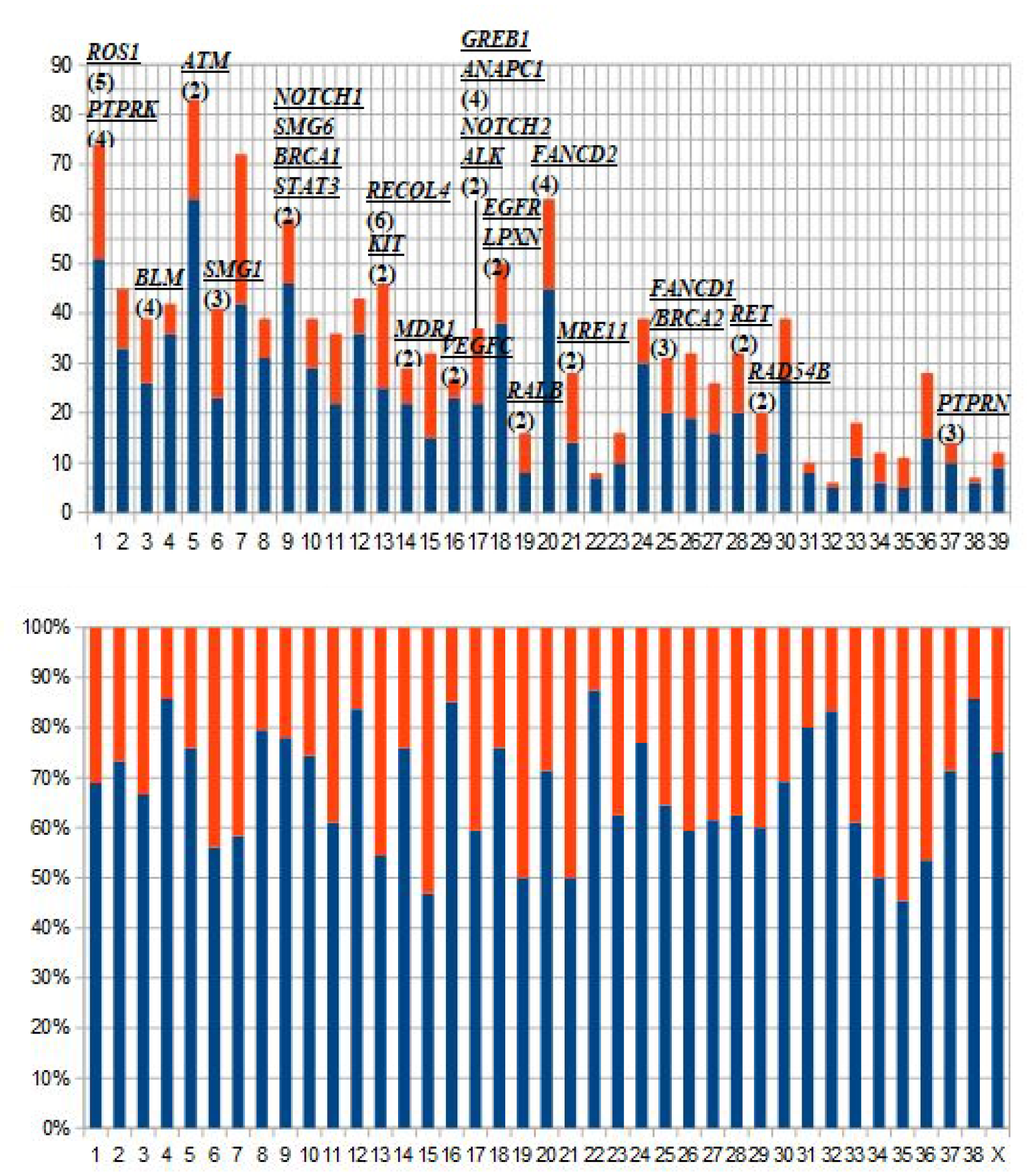
Genome-wide distribution of CTVT multiple-hit candidates of oncological relevance. Upper panel: canine chromosomes representation as vertical bar graph. Vertical bars correspond to each canine chromosome shown on x-axis (1-38 and X; areas corresponding to quantified second-hit and multiple-hit candidates are shown respectively in blue and red; quantified absolute values corresponding to number of identifed candidates representation by bar height show on y-axis to the left; hemizygous X chromosome labelled as 39). Chromosome-wide distrbution of identified multiple-hit candidates of oncological relevance. Individual CTVT loci are imposed above the bar graph and assigned to the area corresponding to specific canine linkage group. The number of hits recored for each mutated candidate is indicated in brackets below the gene name (see Table III for details). Top ranking identification of CTVT-*RECQL4* (corresponding to referrence chr 13) accumulated six mutations. The other relevant identifcations CTVT-*BLM*, CTVT-*FANCD2* and CTVT-*FANCD1/BRCA2* are shown above chr 3, chr 20 and chr 25 respectively. CTVT-*SMG6* (chr 9), CTVT-*KIT*(chr 13), CTVT-ALK (chr 17), CTVT-*NOTCH2* (chr 17) and CTVT-*LPXN* (chr 18) accumulated at least one additional structural variant (see Decker set SV for details) Lower panel: percent stacked values representation of canine chromosomes corresponding to vertical bar graph labelled and presented in upper panel. Chromosome-wide distrbution indicates quantified second-hit candidates expectedly dominate the number of multiple-hit candidates (except for chr 15, 19, 21, 34, 35 and possibly 36 approaching an even distribution).

Due to sourcing issues experienced in accessing the original sets of short-read data deposited by Murchinson and colleagues and preferable data structure, the analyzed data were derived from the set compiled by Decker and colleagues (2015). Hence, conveniently structured Decker set defines a core set used for subsequent manipulations and provides the compendium of variants concordant with two independently sampled aneuploid tumours: 24T and 79T. Since Decker set was derived from the Murchinson sets by the improved filtering of common variants circulating in dog population - we henceforth assumed that – of available genomic short-read samplings, compiled Decker set offers the most representative catalogue of conserved variants acquired, into prototype aneuploid tumour genome, prior to global expansion of canine transmissible venereal sarcoma (CTVT).

### Aneuploid tumour short-read sequencing data acquisition, processing and handling

Data were initially acquired from the variant set compiled by Decker, listing ‘high-quality, high-confidence, putative somatic, protein-changing variants that were found in both tumors’ (Decker et al. 2015). The above dataset was split into 39 individual files, each corresponding to every specific canine linkage group and neglected the unassigned chromosome (‘chrUn’) as indicated in Murchison et al. (2014). Each of 39 individual files was then inspected for the convergence of multiple non-synonymous CTVT variants on the single predicted gene: the mutational pattern indicative for Knudsonian ‘two-hit’ kinetics. CanFam3.1 chromosomal coordinates were maintained over that step, to reflect on the projection of aneuploid sequence onto the structure of the reference canine assembly (RefSeq, haploid-type assembly accession: GCF_000002285.3). Sequential GeneBank accession numbers are given below, including numbers of proteins predicted per chromosome and the total length of assembled chromosomes [values derived from domestic dog assembly available via <https://www.uniprot.org/proteomes/UP000002254>; numbers are given in brackets and followed by total length given in Mb, and RefSeq identifiers corresponding to univalent representation of 38 canine autosomes and singular X chromosome included in reference assembly of female boxer chromosome complement: Chr1 (1548/122,678,785) NC_006583; Chr2 (946/85,426,708) NC_006584; Chr3 (622/91,889,043) NC_006585; Chr4 (690/88,276,631) NC_006586; Chr5 (1264/88,915,250) NC_006587; Chr6 (1116/77,573,801) NC_006588; Chr7 (849/80,974,532) NC_006589; Chr8 (756/74,330,416) NC_006590; Chr9 (1425/61,074,082) NC_006591; Chr10 (826/69,331,447) NC_006592; Chr11 (697/74,389,097) NC_006593; Chr12 (722/72,498,081) NC_006594; Chr13 (485/63,241,923) NC_006595; Chr14 (441/60,966,679) NC_006596; Chr15 (613/64,190,966) NC_006597; Chr16 (762/59,632,846) NC_006598; Chr17 (549/64,289,059) NC_006599; Chr18 (895/55,844,845) NC_006600; Chr19 (245/53,741,614) NC_006601; Chr20 (1141/58,134,056) NC_006602; Chr21 (584/50,858,623) NC_006603; Chr22 (282/61,439,934) NC_006604; Chr23 (416/52,294,480) NC_006605; Chr24 (640/47,698,779) NC_006606; Chr25 (504/51,628,933) NC_006607; Chr26 (608/38,964,690) NC_006608; Chr27 (658/ 45,876,710) NC_006609; Chr28 (481/41,182,112) NC_006610; Chr29 (233/ 41,845,238) NC_006611; Chr30 (514/ 40,214,260) NC_006612; Chr31 (288/39,895,921) NC_006613; Chr32 (269/38,810,281) NC_006614; Chr33 (283/31,377,067) NC_006615; Chr34 (296/42,124,431) NC_006616; Chr35 (247/26,524,999) NC_006617; Chr36 (217/30,810,995) NC_006618; Chr37 (303/30,902,991) NC_006619; Chr38 (253/23,914,537) NC_006620 and ChrX (1013/123,869,142) NC_006621].

### Canine-human synteny, orthology data and relevant morbid information

Synteny information and region comparison were extracted from, and adjusted in, Ensembl_v92-94. Canine-human 1-1 ortholog pairs manipulations were conducted in Ensembl enabled BioMart (<http://www.ensembl.org/biomart/martview/>) specifying the OMIM (An Online Catalog of Human Genes and Genetic Disorders <http://www.omim.org>) identifiers and ‘morbid condition’ as attributes to extract the candidates of oncologic relevance. The regions of canine-human synteny embedding the identified multiple-hit candidates were eventually inspected for the occurrence of CTVT structural variants listed in the Decker set.

### Canine RecQ-like helicase alignment and theoretical modeling

Canine Bloom syndrome RecQ-like helicase ENSCAFP00000043065 (ENSCAFG00000012385 gene on Chr 3:53406975-53506731) and canine RecQ-like helicase 4 ENSCAFP00000002283 (ENSCAFG00000001569 gene on Chr 13:37931119-37937247) were included into a comparative alignments of RecQ-helicase protein sequences derived from following non-canid species: BLM sequences - *H. sapiens* (ENSP00000347232), *F. catus* (ENSFCAP00000029406), *M. musculus* (ENSMUSP00000127995) and RECQL4 sequences - *H. sapiens* (ENSP00000482313), *F. catus* (ENSFCAT00000014395), *M. musculus* (ENSMUSP00000044363), respectively with MAFFT (<https://mafft.cbrc.jp>, with relevant references). CTVT-RecQ varinats derived from Decker set (SNVs) and were incorporated manually into full-leght alignment file and mapped with regard to positions of RecQ variants inferred from non-canid species derived from cited literature (see figure description for sequence identifiers derived from additional species). RecQ protein domain signatures were predicted with PROSITE (<https://prosite.expasy.org>, with relevant references). Based on canine-human alignments, amino-acid sequences of canine RecQ-helicases were folded onto template crystal structures of BLM and RECQL4 catalytic helicase cores from (Swan et al. 2014; Newman et al. 2015; and Kaiser et al. 2017) with SWISS-MODEL protein stucture ExPASy (<https://swissmodel.expasy.org>, with relevant references). The following PDB files (given in supplement section: model_02-4o3m.1.A-CF-BLM-WT.pdb; model_01-4cgz-CF-BLM-WT.pdb; CF-RECQL4-Kaiser2017-model_01.pdb), were manipulated with UCSF Chimera ver. 1.12 (<https://www.cgl.ucsf.edu/chimera/>, with relevant references), to visualize CTVT-RecQ variants derived from Decker set imposed onto canine catalytic helicase core structural models.

## THEORY

Cancer genomes stem out of initially diploid cells through uncontrolled proliferation and acquire a range of structural and numerical variants essentially distracting from euploid chromosomal complement. Cancer genome data presented as sequence aggregates - an alignment of sequence reads over the assembled chromosomes of the reference genome sequence – form a univalent letter-coded string of base sequence rather than the separate assemblies of two haplotypes.

With an aggregation process, alignment of the tumour sequence reads identifies the individual variants (eliminating the sequencing errors and contamination with the wild-type DNA) but the origin of parental chromosomes is often neglected. The parental line of descent of chromosomes is required to correctly assign co-inherited variant alleles for each assembled chromosome – i.e. the phase of the tumour haplotypes (Aguiar et al. 2014). Reconstruction of an aneuploid tumour haplotype phase appears clinically relevant albeit technically challenging (complicated by intratumour variability, copy number aberrations, translocation patterns, genome dosage alterations etc). The relevance of tumour variants haplotype phase is illustrated in cases where assigning of the convergence of multiple non-synonymous tumour variants onto a single gene would test for the Knudsonian two-hit kinetics (Knudson 1971; 2001). In a diploid state, two-hit kinetics assumes the inheritance model where a first deleterious mutational event is followed by additional out-of-phase mutational events. Resulting is a hetero-allelic configuration recapitulating the somatic compound *trans*-heterozygous state; where pathogenic mutational events affected the alleles on both maternal and paternal line of descent.

Accordingly, our objective here was to identify the variant candidates – mutational patterns indicative for two-hit kinetics - tumour genes accumulating the multitude of different alleles. Problematic issue of tumour haplotype phasing is approached here by providing the theoretical estimates: a Table I is given showing the expected rates depending on the intensity of mutational convergence. The values included into a Table I were calculated using the following formula:

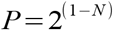

where (*P*)-denotes probabillity of multiple mutations attaining somatic all-*cis* heterozygosity and (*N*)-denotes number of mutational events perceivably affecting the out-of-phase candidate gene.

## RESULTS

Previously reported analyses of mutated Sticker sarcoma genes focused on curation of inconsistencies between the high-quality short-read DNA sequencing data (Murchison et al. 2014 and Decker et al. 2015) and the reference canine genome sequence (Lindblad-Toh et al. 2005). Those efforts have resulted in the unprecedented compendium of tumour genes mutated in CTVT, however - the perceived cases of the somatic compound heterozygosity - have not been assessed systematically. Therefore, we approached the compendium of mutant transmissible tumour genes and browsed for candidate CTVT non-synonymous variants indicative for the perceived hetero-allelic configurations. Provided CTVT genome has acquired many mutations in a number of different genes, to systematically search the mutant variants of canine transmissible sarcoma genes, we mined the structured Decker data-set. Implemented mutation mining strategy was motivated by the multi-mutation theory of cancer (Nordling 1954) followed by the generalization of Knudson ‘two-hit’ hypothesis (Knudson 1971) and assumed the inheritance model where first deleterious mutational event (the first hit) is followed by the additional, somatic mutational events at the remaining wild-type allele (the second hit).

### Genomic landscape of candidate canine transmissible tumour alleles indicative for heteroallelic configuration

The attained genome-wide mining scheme for the generalized ‘two-hit’ inheritance model candidates returned the estimate of 886 doubly mutated candidates and 415 candidates accumulating more than two pathogenic SNV variants concordant in both sampled transmissible tumour isolates. We refer those two initial categories as second-hit candidates and multiple-hit candidates respectively. To evaluate the relative contribution of hypothetical canine linkage-groups to the global landscape of the genomic space occupied by the multiply-mutated out-of-phase candidates we visualized the tumour allele distribution in a chromosome-wide manner, maintaining the WT boxer biased structure of the Decker set CTVT variants (Fig. 1, 2, 3A and 4A).

**Figure 3.**
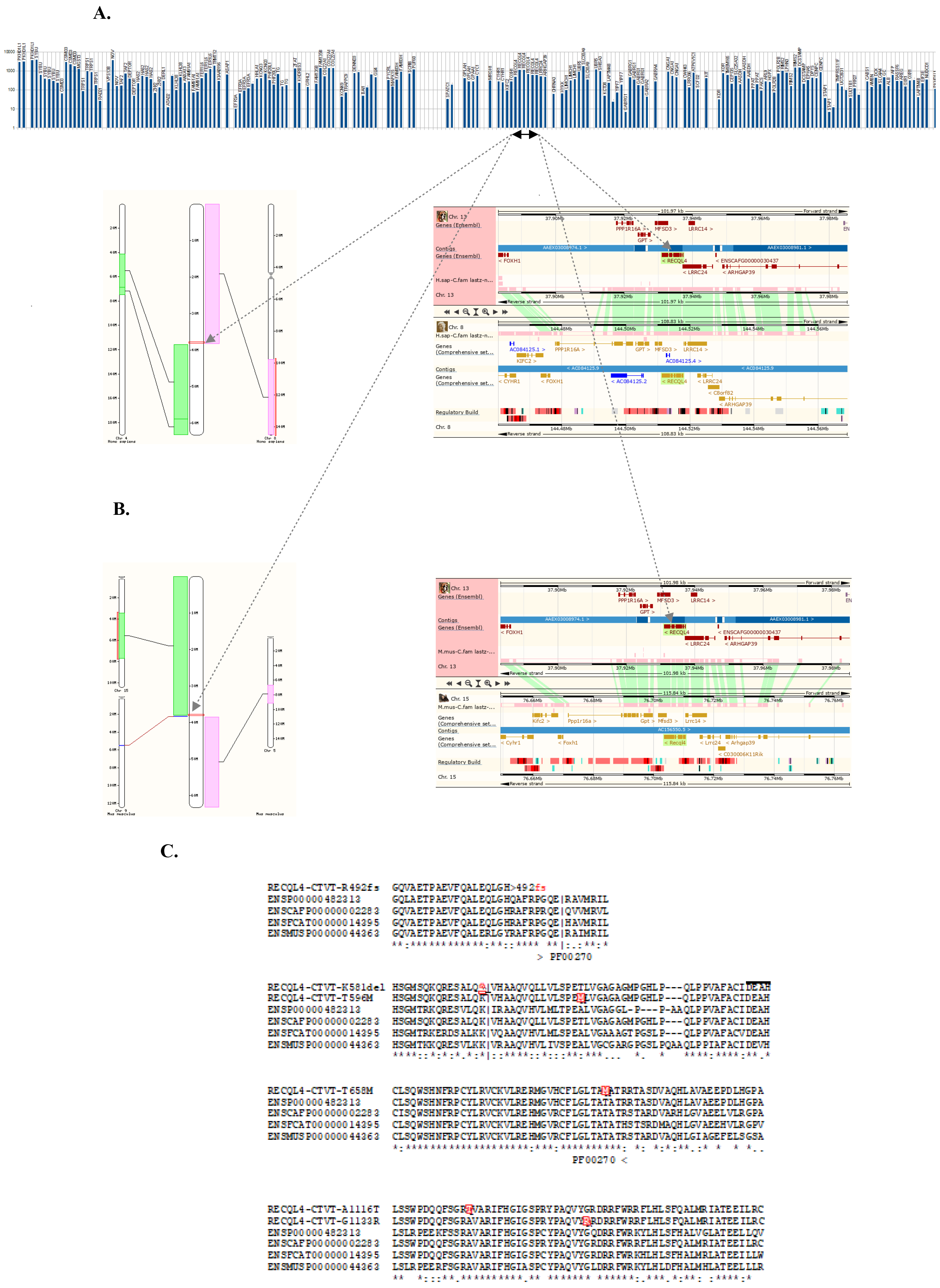
Mutational spectrum of CTVT-RECQL4. **A.** Visualisation of CTVT mutational events aligning with canine chr 13. Chromosome-wide horizontal row stacked bar-graph distribution of SNVs listed with Decker set. Each vertical bar in the top panel represents SNV as a log-scale of the distance (absolute number of residues) of a given non-synonymous substitution from the translational start. (Empty slots correspond to small variants other than strictly SNV i.e. in-frame insertions or deletions). Cluster of six mutational events affecting CTVT-RECQL4 appear in the center (specified by underlined segment). CTVT-RECQL4 cluster is flanked by nearest second-hit candidates: CTVT-CYHR1 and CTVT-ARHGAP39. **B.** Syntenic landscape surrounding multiply-mutated CTVT-RECQL4. Grey arrows connect CTVT-RECQL4 cluster with relevant Ensembl graphics. Chromosomal ideograms shown (left) denote the approximate position of canine *RECQL4* (indicated by red rectangle) with respect to genomic synteny between canine chr 13 and distal tip of the long arm of human chr 8 (pink colored) and mouse chr 15 (green colored) on upper and lower panel respectively. CTVT-*RECQL4* local synteny (right) presented as ~100kb Ensembl window. Gene-chain alignment is indicated by green lines between images, extending on both sides of *RECQL4* (highlighted in green, center of each image). *RECQL4* gene syntax around canine locus (ENSCAFG00000001569) 13:37931119-37937247 is conserved with human *RECQL4* (ENSG00000160957) 8:144511288-144517845 and mouse *Recql4* (ENSMUSG00000033762) 15:76703553-76710548 (upper and lower panel respectively). Note the structural CTVT variants are excluded from above region (see text and supplement table for details and coordinates) suggesting the CTVT gene syntaxt is preserved in genomic region presented. **C.** Amino-acid residue resolution view of alterations to CTVT-RECQL4 concordant in both tumour isolates. Following four mutations expectedly affecting predicted RecQ-like DNA helicase domain are shown in comparative alignment context: frameshift variant RECQL4-CTVT-R492fs (13:3793472137934722), in-frame deletion of lysine codon RECQL4-CTVT-K581del (13:37934024-37934027) removing the splice donor, missense variants RECQL4-CTVT-T596M (13:37933906) and RECQL4-CTVT-T658M (13:37933646). Two mutations altering conserved c-terminal RECQL4 block of amino-acid residues (Bachrati and Hickson 2003) outside of helicase domain: RECQL4-CTVT-A1116T (13:37931370) and RECQL4-CTVT-G1133R (13:37931214). CTVT-RECQL4 altered resiudes are shown in red font. Distinct DEAH box helicase domain motif is indicated by the horizontal line. RECQL4 sequences of following mammalian species are shown in protein alignment: *H. sapiens* ENSP00000482313, *C. familiaris* ENSCAFP00000002283, *F. catus* ENSFCAT00000014395 and *M. musculus* ENSMUSP00000044363. Comparative alignment indicates that CTVT-RECQL4 variants alter conserved and / or invariant residues in aligned vertebrate sequences except for T596M. [Alignment legend: ‘|’ denotes the position of splice donor in the nearest downstream intron; ‘>‘ denotes start and ‘<‘ end of DEAH box helicase domain PF00270/IPR011545 (resiudes 496-663), overlapping with Prosite profile prediction (residues 502677) Helicase superfamily 1/2, ATP binding domain, ‘<u>^</u>’ denotes the K581 in-frame deletion]

**Figure 4.**
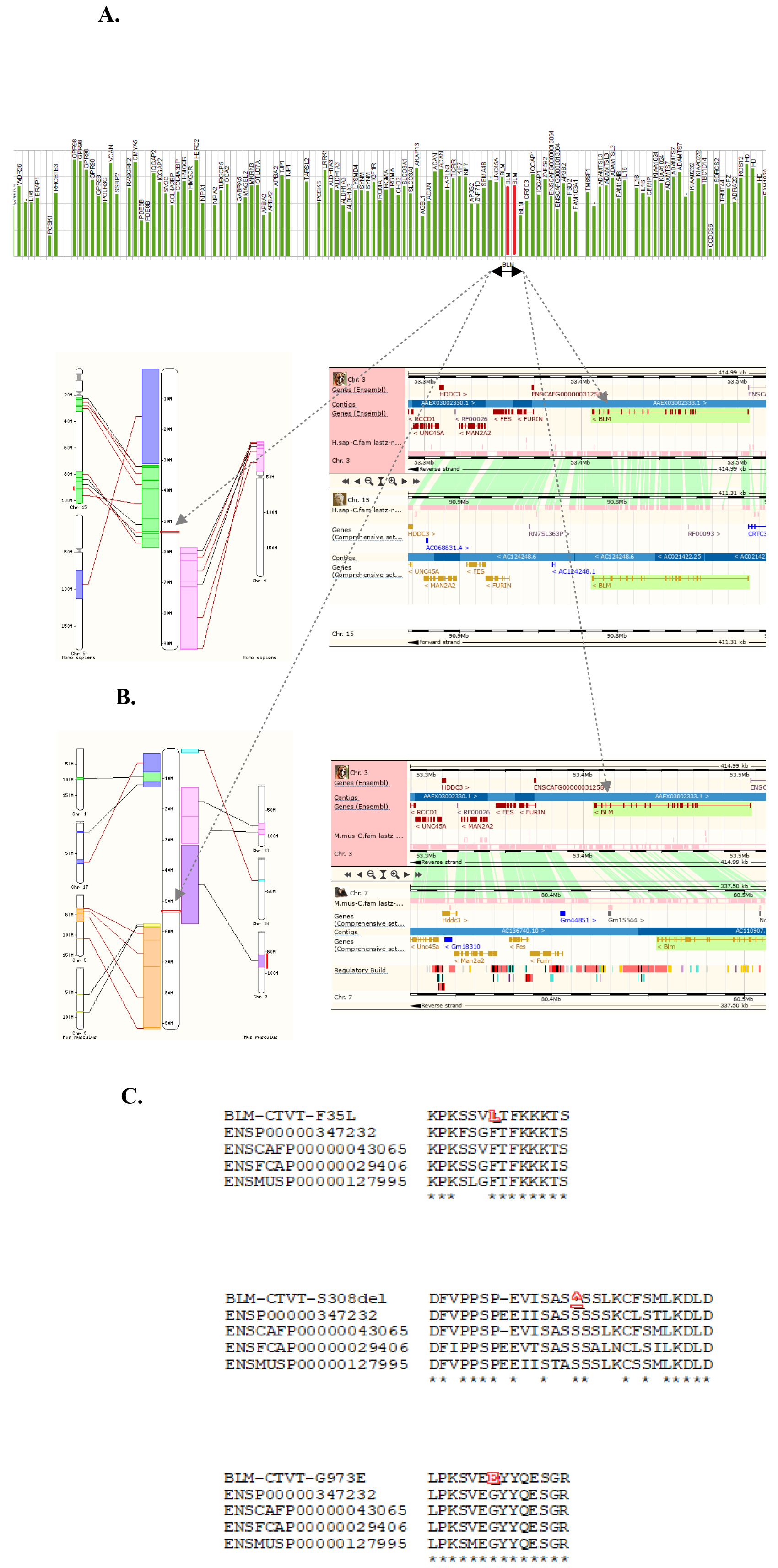
Mutational spectrum of CTVT-BLM. **A.** Visualisation of CTVT mutational events aligning with canine chr 3. Chromosome-wide horizontal row stacked bar-graph distribution of SNVs listed with Decker set. Each vertical bar in the top panel represents SNV as a log-scale of the distance (absolute number of residues) of a given non-synonymous substitution from the translational start. Two empty slots between BLM-CTVT-F35L and BLM-CTVT-G973E, corresponding to small variants other than strictly SNV i.e. in-frame deletion of S307-308 were replaced with adequate red vertical bars. Cluster of four (three unique, see Decker set for details) mutational events affecting CTVT-BLM appear in the center (specified by underlined segment). CTVT-BLM cluster is flanked by nearest second-hit candidates: CTVT-KIF7 and CTVT-IQGAP1. **B.** Syntenic landscape surrounding multiply-mutated CTVT-BLM. Grey arrows connect CTVT-BLM cluster with relevant Ensembl graphics. Chromosomal ideograms shown (left) denote the approximate position of canine *BLM* (indicated by red rectangle) with respect to genomic synteny between canine chr 3 and distal tip of the long arm of human chr 15 (green colored) and mouse chr 7 (purple colored) on upper and lower panel respectively. CTVT-*BLM* local synteny (right) presented as ~400kb Ensembl window. Gene-chain alignment is indicated by green lines between images, extending on both sides of *BLM* (highlighted in green, center of each image) except for ENSCAFG00000031258. *BLM* gene syntax around canine locus (ENSCAFG00000012385) 3:53406975-53506731 is conserved with human *BLM* (ENSG00000197299) 15:9071732790816165 and mouse *Blm* (ENSMUSG00000030528) 7:80454733-80535119 (upper and lower panel respectively). Markedly *FES* gene (15q26.1) used originally in *BLM* linkage mapping (German et al. 1994, Ellis et al. 1994) as well as adjacent *FUR* gene (Ellis et al. 1995, Ellis & German 1996), appear in presented corresponding gene-chain alignments of canine and mouse, consistent with preserved gene syntax. Note the compared regions includes reverse strand of canine chr 3 and forward strand of human chr 15. The structural CTVT variants are excluded from above region (see text and supplement table for details and coordinates) suggesting the CTVT gene syntaxt is preserved in genomic region presented. **C.** Amino-acid residue resolution view of alterations to CTVT-BLM concordant in both tumour isolates. Following mutations affecting predicted canine BLM RecQ-like DNA helicase are shown in comparative alignment context: BLM-CTVT-F35L (3:53471822), BLM-CTVT-S307-308del (3:53469883-53469886) and BLM-CTVT-G973E (3:53433860). CTVT-BLM altered resiudes are shown in red font. BLM sequences of following mammalian species are shown in protein alignment: *H. sapiens* ENSP00000347232, *C. familiaris* ENSCAFP00000043065, *F. catus* ENSFCAP00000029406 and *M. musculus* ENSMUSP00000127995. Comparative alignment indicates that CTVT-BLM variants alter conserved and / or invariant residues in aligned vertebrate sequences except for BLM-CTVT-S307-308del (‘<u>^</u>’ denotes the in-frame deletion) where inferring on relevance of the contraction of oligo-serine stretch is problematic.

Chromosome-wide prevalence of the multiply-mutated candidates demonstrated, that unlike the hemizygous CTVT X chromosome, candidate distribution was expectedly uniform when compared the relative contribution across autosomal loci i.e. the regions corresponding to largest canine autosomes have contributed proportionally more candidates compared with hemizygous X. In absolute values, CTVT hemizygous X chromosome (of a size and coding capacity comparable with largest canine autosomes) have only contributed 9 doubly mutated and 3 multiply-mutated candidate genes - contribution comparable to values recorded for smallest canine autosomes. This trend is reflected by Pearson correlation coefficient scores indicating strong (second-hit candidates; PCC 0.8805-0.8257, significant at *P*-value < .00001 - .000016 respectively) to moderate (multiple-hit candidates; PCC 0.617-0.5598, significant at *P*-value .000037 - .000211 respectively) positive correlation between the chromosome-wide distribution of multiply-mutated candidates and the number of autosomally encoded proteins (Table II, and Methods section for source values). Hemizygous CTVT X chromosome in aneuploid tumour complement, however, encountered the complex pattern of inter-chromosomal rearrangements and the number of X derived fragments have translocated to other genomic locations, hence certain regions of CTVT X chromosome might deviate from the expected copy number.

The final data-mined set included at least one exceptionally informative candidate, where the convergence of variants seems to support Knudsonian two-hit kinetics model. The remarkable example of convergence concerns a multiply-mutated CTVT gene encoding for TLR5 (enSCAFG00000011368) on Chr 38; the case convergence of two, biallelically exclusive, pathogenic variants onto a single codon was uncovered in Decker set. CTVT-*TLR5* converged both stop-gaining (ochre) and frame-altering SNV affecting Y847 encoded by *tac* codon (38:23704571-23704573; Table III; further details included in Decker set). Exceptional convergence of two mutually exclusive CTVT variations onto a single codon implies out-of-phase variants constellation and is consistent with the *trans*-heteroallelic model. Thereof, assuming broadly autosomal CTVT 2n copy number, detection of two allelically exclusive variants (annotated in aggregated unphased sequence consensus), could be regarded as a *bona fide* example of Knudsonian two-hit kinetics.

### Detection of gene-size bias among multiple-hit out-of-phase candidates

To evaluate the mutational space occupied by the CTVT compound heterozygous candidates we data-mined the 415 multiple-hit out-of-phase candidates accumulating more than two deleterious or pathogenic SNV variants concordant in both sampled transmissible tumour isolates. The results of the chromosome-wide mapping of CTVT short-reads aligned to the reference canine assembly have specified the subset of 51 loci representing examples of the most intensely mutated Sticker sarcoma genes represented in the Decker set. The results of chromosome-wide mapping (summarized in Fig. 1 and Table III) demonstrate the most intensely mutated multiple-hit candidates have accumulated between three and nine mutations per CTVT encoded tumour gene. The genome-wide subset of most intensely mutated multiple-hit candidates however could be biased towards loci encoded by genes occupying the extremely large chromosomal space or encoding for exceptionally large proteins. Consistent with this perspective we noted that some the most intensely mutated multiple-hit candidates represent the extreme examples. The extreme examples of tumour encoded mutated multiple-hit candidates include *LRP1B* and *TTN* (Table III). CTVT encoded *LRP1B* accumulated 7 variants across gene sized at 1,29Mb and *TTN* accumulated 8 variants across protein coding sequence sized at 105,54Kb. Given the two above extreme examples listed in Table III and other examples of multiple-hit CTVT candidates not listed here, but present in the Decker set e.g. canine X-linked *DMD* gene (not shown; McGreevy et al. 2015 and Kornegay 2017) encoded by the gene sized at ~2Mb, the perceived gene-size bias was addressed in more systematic terms. Specifically, based on reference canine assembly providing gene size and protein coding sequence estimates for predicted canine genes, the relative mutational load was estimated converting the absolute numbers, corresponding to CTVT mutations acquired in multitude into tumour genes, to local mutational frequency rates. This analysis have established that some of the most intensely mutated CTVT genes listed in Table III, are characterized by low local non-synonymous SNV frequency rates. Variants across extremely sized genes are estimated at range of >10 non-synonymous SNV per Mb of overall gene sequences e.g. local mutational frequency rate of *LRP1B* gene is estimated at rate of 5,43 non-synonymous SNV per Mb of gene sequence. Variants across genes encoding for exceptionally large proteins coding sequence are estimated at range of >0,5 non-synonymous SNV per Kb of protein coding sequence e.g. local mutational frequency rate of TTN protein coding sequence is estimated at rate of 0,08 non-synonymous SNV per Kb. Interestingly this analysis have revealed the other extreme of the gene-size bias: CTVT encoded multiple-hit out-of-phase candidates characterized by the exceptional theoretical intensity of local non-synonymous SNV rates. For example, local non-synonymous SNV frequency rates estimated for predicted canine gene *MAFF* (ENSCAFG00000029842) appear at 500 non-synonymous SNV per Mb of gene sequence and 10 non-synonymous SNV per Kb of protein coding sequence.

### Mutation mining of multiple-hit out-of-phase candidates of relevance in oncology

Having established the mutational space of the extreme cases of CTVT multiple-hit candidates, the further analyses were focused on non-synonymous variants affecting the genes of oncologic relevance. The results of the chromosome-wide mapping of CTVT short-reads aligned to the reference canine assembly have specified the subset of 30 loci where multiple local tumour specific non-synonymous variants affected the canine genes of perceived oncologic relevance. The results of chromosome-wide mapping (summarized in Table IV and V) demonstrate the subset of CTVT encoded genes of oncologic relevance have accumulated between two (second-hit candidates) and six (multiple-hit candidates) non-synonymous SNVs per tumour gene. Since the theoretical expectations (Table I) predict the second-hit out-of-phase candidates (Table IV) retain only 50% likelihood of attaining the *trans*-heterozygous Knudsonian inheritance model at 2n copy number, subsequent analyses involved the subset of tumour genes referred as multiple-hit out-of-phase candidates (Table V). Multiple-hit out-of-phase candidates retain equal or greater than 75% likelihood of attaining the *trans*-heterozygous Knudsonian inheritance model at broadly assumed CTVT autosomal 2n copy number state (Murchison et al. 2014). This step had enabled us to focus on strongest somatic compound heterozygosity candidates encoded by the aneuploid transmissible canine tumour. Based on applied canine-human gene orthology criteria, by means established in comparative oncology (Table VI), four CTVT multiple-hit candidates were identified as encoding for orthologs of relevant human hereditary neoplastic syndromes (Table VII). Amongst four identified CTVT multiple-hit out-of-phase candidates two encoded for RecQ-like helicases: RECQL4 and BLM. Two other identified candidate multiple-hit canine factors encoded by aneuploid CTVT listed in Table VII are described as FANCD1 and FANCD2. Notably, four final multiple-hit CTVT candidates are classified as morbid conditions referred as DNA-repair deficiency disorders (relevant MIM gene description and morbid accessions are given in Table VII). We note the above human maladies, referred as hereditary neoplastic syndromes, represent recognized chromosome instability syndromes (Taylor 2001) have not been described in domestic dogs. Moreover, we note that in previous reports concerning the CTVT mutational load analyses (Murchison et al. 2014) those factors of oncologic relevance were unnoticed with the exception of FANCD1 described in Decker et al. (2015) under the alternative symbol (BRCA2) maintained by the HUGO Gene Nomenclature Committee. Provided the description focus on the canine RecQ-like helicases (were FANC axis is regarded as auxiliary pathway) we prefer to refer canine ortholog as FANCD1/BRCA2.

**Table VII.**
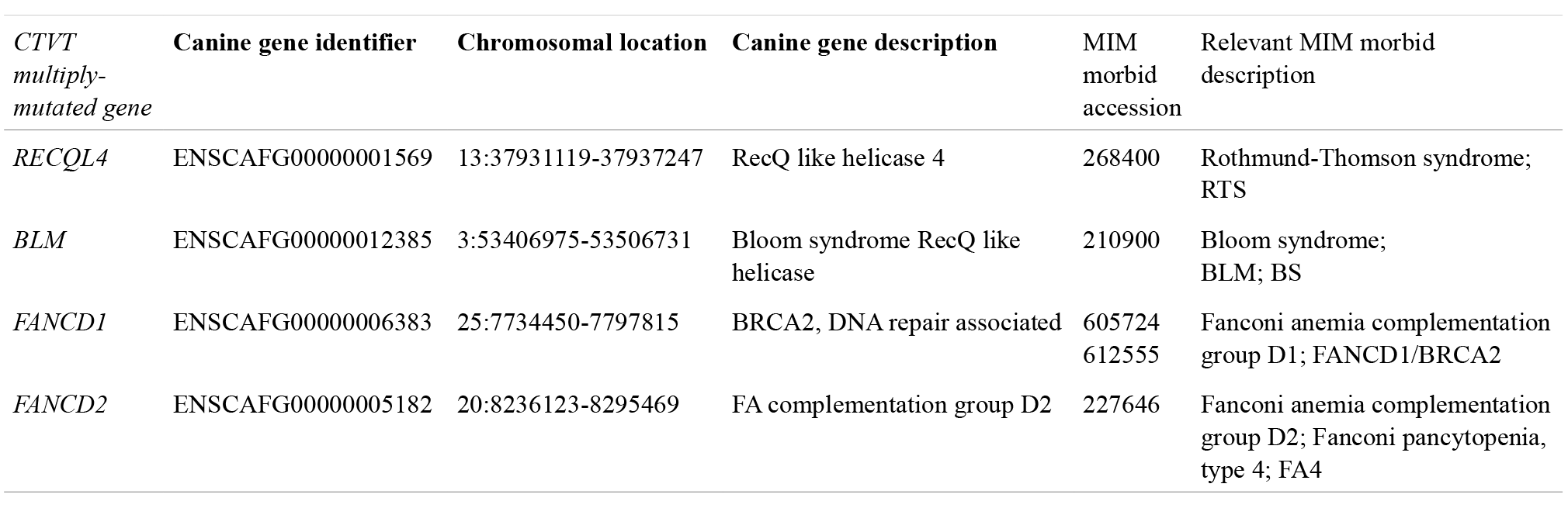
CTVT multiply-mutated genes encoding for homologs of relevant human hereditary neoplastic syndromes.

### Two predicted canine RecQ-like helicases represent multiple-hit out-of-phase candidates encoded by the aneuploid transmissible tumour

Previous analytical steps have established that two multiple-hit out-of-phase candidates present in subsets of the most intensely mutated Sticker sarcoma genes encode for predicted canine RecQ-like helicases. Canine RecQ-like helicases, however, in contrast to human syndrome causing RecQ helicases strongly implicated in cancer predisposing heritable syndromes (reviewed in Bjergbaek et al. 2002, Bachrati & Hickson 2003), have not been directly characterized clinically either genetically. By means established elsewhere in comparative oncology, based on canine-human gene homology criteria, the applied multiple-hit mining scheme have specified transmissible tumour encodes two RecQ-like helicases affected by multiple SNV non-synonymous polymorphisms: RECQL4 and BLM. A transmissible tumour encoded canine RECQL4, in particular, have accumulated six different non-synonymous SNV polymorphisms. Based on the additional systematic measures applied, the local non-synonymous SNV frequency rates estimated (Table V) CTVT-RECQL4 have accumulated considerably more than 600 non-synonymous SNV per Mb of gene sequence (the precise estimate is 978,79 non-synonymous SNV per Mb, given the predicted canine gene encoding for RECQL4 is sized at 6,13 Kb). The exceptional rate of local non-synonymous SNV accumulated by CTVT-RECQL4 indicates a transmissible tumour encoded RECQL4 helicase represents the most intensely mutated multiple-hit out-of-phase candidate of oncological relevance detected by the applied standards. Based on the absolute values (Table I), theoretical expectations specify that assuming autosomal 2n copy number state, cumulatively CTVT-RECQL4 retain 96,87% likelihood of attaining the compound *trans*-heterozygous Knudsonian inheritance model consistent with out-of-phase configuration. Conversely, operating at the above 2n copy number assumption, same expectations specify the chance of retaining the intact wild-type CTVT-RECQL4 allele is 3,13% consistent with all-*cis*-heterozygous allele configuration. For comparison, the other identified helicase, CTVT-BLM is mutated at the rate comparable to FANCD1/FANCD2 (Table V) with mutational rates typical for other ‘reasonably sized’ multiple-hit out-of-phase candidates presented.

### Comparative analysis of multiple-hit out-of-phase candidates CTVT-RECQL4 and CTVT-BLM extends the association with human genes causally implicated in progeroid, cancer predisposing heritable syndromes

Human RECQL4 (Rothmund-Thomson syndrome helicase) and BLM (Bloom syndrome helicase) are often classified together with WRN (Werner syndrome helicase) as heritable cancer predisposing RecQ repair-deficiency progeroid syndromes and differentially diagnosed with other chromosome instability syndromes including Ataxia-telangiectasia (ATM), and Fanconi anemia (FANC) (Taylor 2001, Amor-Gueret 2004, Wang & Plon 2016, Larizza et al. 2010). Since identified canine multiple-hit candidates encoding CTVT-RECQL4 and CTVT-BLM have affected two different ATP-dependent RecQ-like helicases, the entire Decker set was re-searched. Those results have confirmed that unlike CTVT-*RECQL4* and CTVT-*BLM*, canine *WRN* (ENSCAFG00000006410) has not accumulated any SNV polymorphisms concordant in two sampled CTVT isolates.

The identification of two CTVT multiply-mutated canine RecQ-like helicases was based on the analysis of SNV polymorphisms; Decker et al. (2015) however included structural variant (SV) analysis in addition to polymorphic SNVs. Structural variants affecting tumour cells can contribute the equivalent of the second-hit and efficiently attain the somatic compound *trans*-heterozygous Knudsonian inheritance model. To evaluate the possible SV contribution to mutational patterns affecting the CTVT encoded RecQ-like multiple-hit candidates, the syntactic criteria were used to independently verify the conservation of the CTVT gene order, and thereof ensure the overall correctness of the canine-human homology-based predictions. First, chromosomal regions predicted to encode CTVT multiple-hit RecQ-like out-of-phase candidates structured in the Decker set as CTVT-Boxer consensus were imposed on Ensembl maps of canine-human synteny. The results of mapping of regions predicted to encode CTVT multiple-hit RecQ-like out-of-phase candidates, structured in the Decker set as CTVT-Boxer consensus, are presented in Figure: 3 and 4 (panels A. and B. respectively for CTVT-*RECQL4* and CTVT-*BLM*). Presented results confirm canine RECQL4 is located in a region of preserved gene syntax both in canine-human RECQL4 (midregion of canine chr 13 and distal end of a long arm of human chr 8q24.3, Kitao et al. 1998; 1999) but also canine-mouse gene-chain alignment (mouse chr 15; given Hoki et al. 2003 and Mann et al. 2005, mouse Recql4 is relevant in further considerations). The parallel results confirm canine BLM is located in a region of preserved gene syntax both in canine-human BLM (mid-region of canine chr 3 and distal end of a long arm human chr 15q26.1, Ellis et al. 1995) but also canine-mouse gene-chain alignment (mouse chr 7; see Chester et al. 1998 for mouse Blm). Second, given the highly aneuploid chromosomal complement of CTVT cells we reasoned the above syntenic alignment could have been interrupted when aligned with the Decker set CTVT-Boxer consensus; thus the transmissible tumour SV described in (Decker et al. 2015) were imposed onto existing map. Two types of CTVT structural variants (SVs) are listed as separate dataset in Decker et al. (2015) : inter- and intra-chromosomal. The results of mapping of the CTVT SVs onto syntenic alignment are included in Table VIII. The results of mapping of the transmissible tumour SVs onto regions encoding CTVT-RECQL4 and CTVT-BLM demonstrate the nearest flanking inter- and intra-chromosomal SV (including copy number variants, CNV) are located at the considerable distance (at least few hundred Kb apart, Table VIII) from multiply-mutated RecQ genes. Collectively, this allowed us to conclude the SV concordant in both sampled transmissible tumour isolates did not affected the multiply-mutated RecQ-like helicases in the prototype CTVT. Consistently this suggests the detected SV have not affected (i.e. neither inter-nor intra-chromosomal rearrangements contributed the equivalent of the second-hit) the multiply-mutated RecQ-like helicases in an early history of CTVT.

### Multiply-mutated CTVT-*BLM* encodes a putative catalytically inactive variant of RecQ-like helicase

Previous steps established two CTVT multiply-mutated canine RecQ-like helicases are regarded as orthologs of human RECQL4 and BLM (Fig. 3 and 4). Hence, predicted as mutually paralogous, multiple-hit candidate tumour genes: CTVT-*RECQL4* and CTVT-*BLM*, are regarded as related to distinct human ATP-dependent RecQ-like DNA helicase family members - classified as loci implicated in the heritable cancer predisposing repair-deficiency progeroid syndromes. Bloom syndrome, in particular, is characterized by strong predisposition to variety of neoplastic conditions of a recessive mode of inheritance. Homozygous or compound heterozygous *BLM* mutation carriers develop a malignant neoplasia of a wide variety of anatomical sites and types occurring at excessive frequency and arising at earlier than typical ages. This cancer predisposition results in various cellular types of neoplasms including hematopoietic, lymphoid, epithelial and other malignant forms. *BLM* encodes Bloom syndrome ATP-dependent RecQ like helicase (Ellis et al. 1995). Bloom syndrome (BS) was originally described as ‘The syndrome of congenital telangiectatic erythema and stunted growth’ (Bloom 1966) and is clinically regarded as pre-malignant state. BS and mutant BLM encoding RecQ like helicase human gene was studied by others (e.g. German et al. 1974; reviewed Ellis & German 1996; German 2004; German et al. 2007; Cunniff et al. 2017) resulting in detailed functional and structural description exceeding by far studies concerning the other relevant helicase RECQL4 (Kitao et al. 1998; 1999). BS is referred as ‘a Mendelian prototype of somatic mutational disease’ (German 1993). Given that notion of German – in a comparative oncology perspective - multiply-mutated CTVT-BLM appears relevant in considerations pertaining to the accumulation of somatic mutations in canine transmissible sarcoma. Heritable ‘Bloom-like syndrome’, however, has not been described in dogs - canine *BLM* (ENSCAFG00000012385) exists solely as a conceptual homology-inferred database prediction. ENSCAFG00000012385 was originally identified as *Cfam-BLM* with reciprocal orthology search (not shown), between species other than man: mouse *Blm* (Chester et al. 1998) and *C. elegans him-6* (Hodgkin et al. 1979; Wicky et al. 2004). CTVT-*BLM* identified here, as transmissible tumour mutant gene affected by the multiple SNVs, was used as a positive control case across the CTVT multiple-hit out-of-phase candidate mining scheme described herein.

CTVT-BLM is mutated at a rate comparable to other multiple-hit out-of-phase candidates, but at the rate lower than CTVT-RECQL4 (Table V). Moreover, the severity, of mutations recorded in CTVT-RECQL4, is perceivably higher than what observed in CTVT-BLM (Fig. 3C and 4C). Specifically, mutational spectrum of CTVT-BLM predicts transmissible tumour encodes allelic variant of a mature protein of length comparable to wild-type *Cfam*-BLM, affected by missense substitutions and inframe deletion - hence, unlikely resulting in a premature protein-translation termination. The evaluation of the perceived cumulative pathogenicity of those variants, however, is expectedly problematic based on the unphased sequence information alone. *BLM*-type family of RecQ-like DNA helicases encode multi-domain proteins; therefore initially the CTVT-*BLM* mutations were categorized by assignment to predicted protein domains. This categorization established one of the multiply-mutated CTVT-BLM substitution variants, G973E, alters the RecQ helicase C-terminal domain (RecQ-Ct, PS51194 - corresponding to residues 879 – 1026 in *Cfam*-BLM). RecQ-Ct region is essential for BLM activity and consequently is essential for the maintenance of genomic stability (Yankiwski et al. 2001). Further, it has been suggested that missense substitutions or frame-preserving deletions in the C-terminal part of BLM are sufficient to cause an increase in a level of chromosome abnormalities. This notion was extended by Guo et al. (2005) suggesting that some missense mutations acquired into RecQ-Ct BLM can lead to the destabilization of the genome and ultimately lead to tumorigenesis.

Consistently, German et al. (2007) have listed a subclass of amino-acid substitutions to RecQ-Ct regarded as Bloom syndrome causing missense alleles – pathogenic variants known to occur in a defined ‘missense-sensitive region’ of BLM helicase. Collectively the registry of Bloom syndrome causing missense alleles and protein domain based resolution analysis enabled mapping of the CTVT-BLM substitution variant G973E. The canine-human RecQ-Ct(PS51194) protein domain alignment uncovered CTVT-BLM G973E maps onto missense-sensitive interval, between tightly spaced G952V and H963Y towards the N-terminus (Fig. 5B and C) and C1055G/S/R towards C-terminus (not shown) of *Cfam*-BLM respectively. To improve the mapping to subdomain resolution, based on the above protein alignment, the predicted *Cfam-BLM* amino-acid sequence was folded onto the template crystal structure of the catalytic helicase core of human BLM (Swan et al. 2014 and Newman et al. 2015). Based on the structure model, CTVT-BLM G973E was found to alter the conserved alpha-helix VE**<u>G</u>**YYQESGR into VE**<u>E</u>**YYQESGR (Fig. 5 and 6) embedded in canine RecQ-Ct(PS51194). The structural framework provided by the theoretical modeling further established the conserved alpha-helix altered by CTVT-BLM G973E is placed in the second lobe of *Cfam-BLM* and constitutes a part of D2 RecA-like domain. D2 along with first RecA-like domain D1 provides the catalytic helicase core with adenosine triphosphate (ATP)-dependent motor activity required for functions of DNA unwinding (Swan et al. 2014 and Newman et al. 2015, reviewed in Cunniff et al. 2017). Bloom syndrome causing, pathogenic missense mutations, however, occur at the conserved amino-acid residues of the catalytic core helicase region of BLM. The D2 embedded, conserved alpha-helix altered by CTVT-BLM G973E, is identical in canine-human sequence alignment. Therefore the corresponding region of canine-human catalytic helicase core (motif VI) consensus was inspected in additional species (Fig. 5C). The multiple protein alignment, of the region embedding the CTVT-BLM G973E altered alpha-helix, specified the particular di-tyrosine adjacent glycine represents an essentially invariant residue (Fig. 5C) even in distantly related invertebrate species.

**Figure 5.**
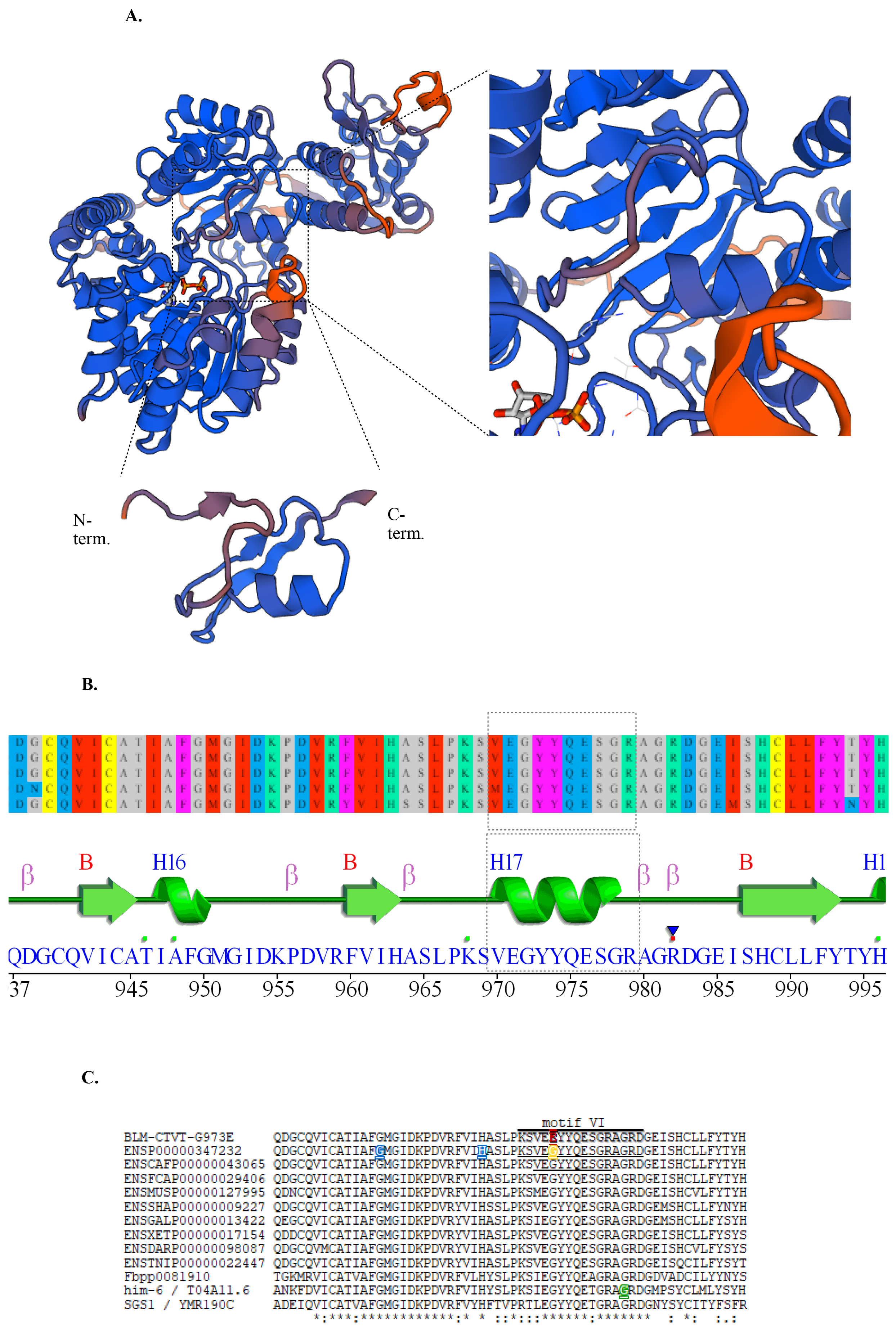
Detailed view of conserved α-helix VE**<u>G</u>**YYQESGR, embedded in the helicase motif VI, altered by CTVT-G973E. ADP moiety shown in ball-stick format. **A.** Swiss-model graphics: catalytic DNA helicase core *Cfam*-BLM (left) rotated with DNA binding region pointing up-right; horizontally oriented conserved α-helix in a center of boxed area. Close-up view of conserved α-helix region (right) in the context of *Cfam*-BLM catalytic DNA helicase core model. Isolated region surrounding conserved α-helix embedded in helicase motif VI (below). **B.** PdbSum secondary structure plot of human Blm PDB ID: 4CGZ (Newman et al. 2015). Linear ideogram of secondary structures corresponding to ca. 60 residues of missense-sensitive region surrounding conserved α-helix (indicated by the grey box). Note theoretical models predictions outside of conserved α-helix embedded in helicase motif VI are sligthly different compared to model by Swan et al. (2014). **C.** Location of missense BLM variants: MAFFT comparative alignment of the above missense-sensitive region in a ClustalW format. CTVT-G973E altering the conserved α-helix is shown in red. Helicase motif VI indicated by horizontal line (and underlined in human BLM sequence). Conserved α-helix is underlined in *Cfam*-BLM. Bloom syndrome causing missense substitutions G952V and H963Y are shown in blue in human sequence. BLM RecQ helicase missense variant G972V that is not currently associated with BS (Mirzaei & Schmidt 2004 and Shastri & Schmidt 2016) is shown in yellow, *him-6* allele (*e1104*) from Hodgkin et al. (1979) altering helicase motif VI G561Q missesnse variant (Wicky et al. 2004) shown in green.

**Figure 6.**
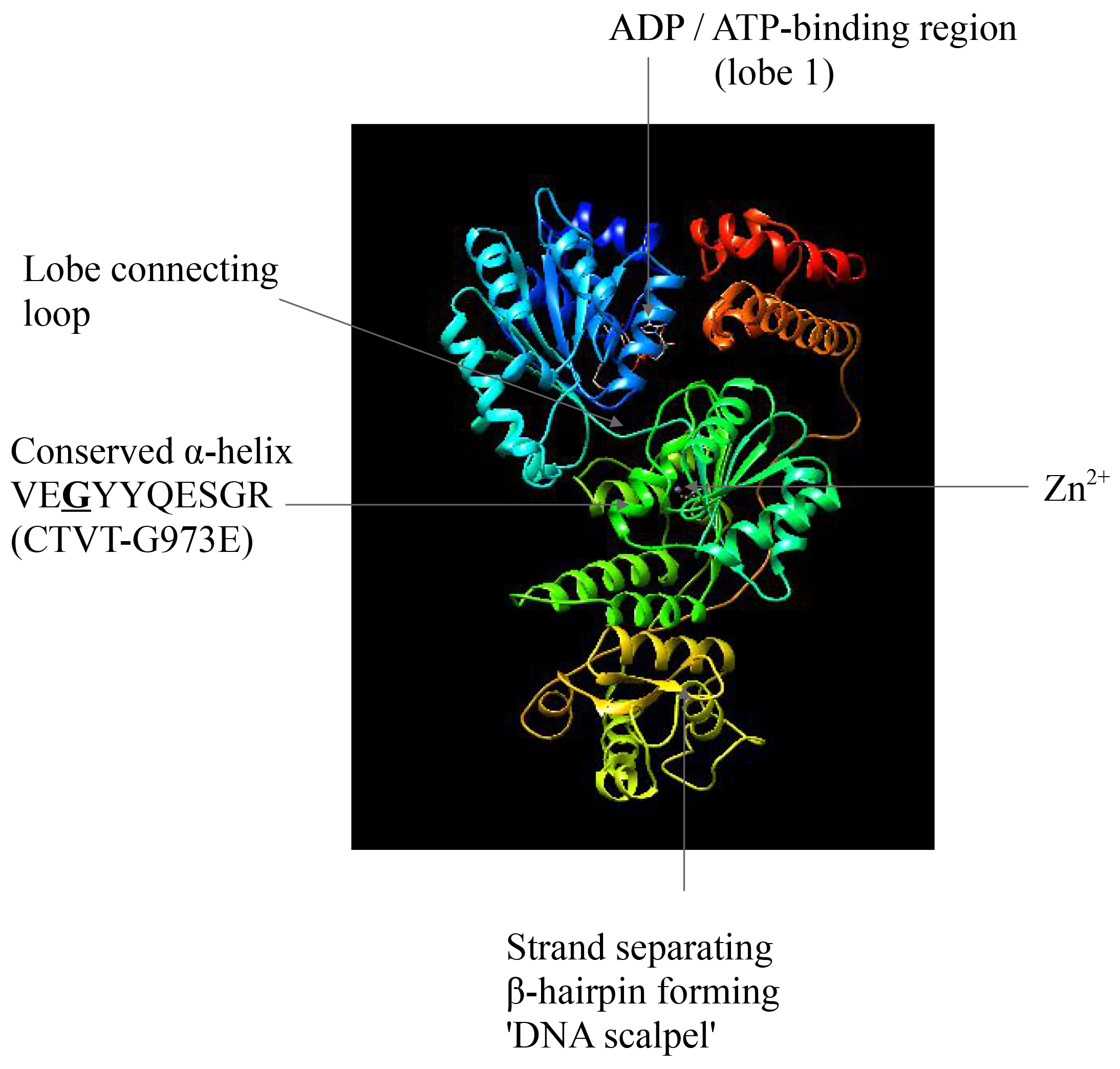
Screen-snipp of interactive model of catalytic DNA helicase core *Cfam*-BLM (ENSCAFG00000012385) folded onto the template crystal structure of the catalytic helicase core of human BLM (Swan et al. 2014) PDB: 4O3M. ADP moiety shown in ball-stick format. Zinc ion shown as blue sphere. DNA omitted for clarity, insead ‘DNA scalpel’ (Cunniff et al. 2017) is indicated by arrow. Conserved α-helix VE**<u>G</u>**YYQESGR, embedded in the helicase motif VI, altered by CTVT-G973E located in second lobe (arrow points to the position of di-tyrosine adjacent glycine). Image generated and colored with standard N-term. →-C-term. code with UCSF Chimera ver. 1.12, demonstrating bi-lobal structure of catalytic DNA helicase core.

Broad phylogenetic representation of the multiple protein alignment of the region embedding the CTVT-BLM G973E altered alpha-helix allowed to further impose the functionally characterized BLM missense alleles derived from diverse range of species. Provided the studies in mice have concentrated on structural *Blm* variations or other types of engineered alleles (i.e. conditional) the missense alleles characterized in genetic model organisms were explored. Namely, *DmBlm* allele (*mus309*) carrying missense E866K (McVey et al. 2007) in a fruit fly and the nematode *him-6* mutant line CB1138 (Hodgkin et al. 1979) carrying homozygous G561Q missense allele (*e1104*) altering the catalytic RecQ helicase region (Wicky et al. 2004). The analysis of the comparative alignment have established that in contrast to *DmBlm* missense mutation *mus309* (aligned in the N-terminal region of DmBlm catalytic RecQ helicase and altering the conserved helicase DEAH motif II) the *him-6* altering missense mutation G561Q aligned in the immediate vicinity of the region embedding the G973E altered alpha-helix of CTVT-BLM (Fig. 5C).

To further improve the mapping resolution, based on extended functional screen for BLM RecQ helicase variants that are not currently associated with BS (Mirzaei & Schmidt 2004 and Shastri & Schmidt 2016), the additional missense loss-of-function alleles were incorporated into subdomain analysis. This step have focused on BLM missense variant G972V (Fig. 5C). According to the comparative alignment, CTVT-BLM G973 corresponds to G972 of BLM (given the presence of inframe deletion in CTVT-BLM listed with Decker set at 307-308 position, resulting in the removal of single residue from tetra-serine repeat) thereof represents the equivalent change to the conserved alpha-helix of CTVT-BLM altered by G973E. Provided G972 is an invariant residue, located in the catalytic helicase core ATP interacting motif VI, Mirzaei & Schmidt (2004) suggested this loss-of-function missense variant is expected to impair coordination between lobe 2 and ATP binding in lobe 1. Given that notion, the equivalent G973 polymorphism in CTVT-BLM, converting invariant glycine to charged residue (altering the conserved helicase motif VI alpha-helix, critically involved in the ATP-dependent helicase activity), is therefore expected to represent missense SNV which fail to efficiently couple the ATP hydrolysis to DNA binding and unwinding. Together the above efforts of theoretical modeling and mapping at the amino-acid resolution of the G973E variant prompted us to conclude that the multiple-hit out-of-phase candidate CTVT-*BLM* encodes a putative catalytic-null.

In parallel, we launched the modeling effort and utilized *Cfam*-RECQL4 amino-acid sequence folded onto the template crystal structure of the segment catalytic helicase core of human RECQL4 (Kaiser et al. 2017). The corresponding model is shown in (Fig. 7) and includes the corresponding position of unphased alleles recorded in multiply-mutated CTVT-RECQL4 (Fig. 3C). Again, the evaluation of the perceived cumulative pathogenicity of above CTVT-RECQL4 variants, is expectedly problematic based on the unphased sequence information alone.

**Figure 7.**
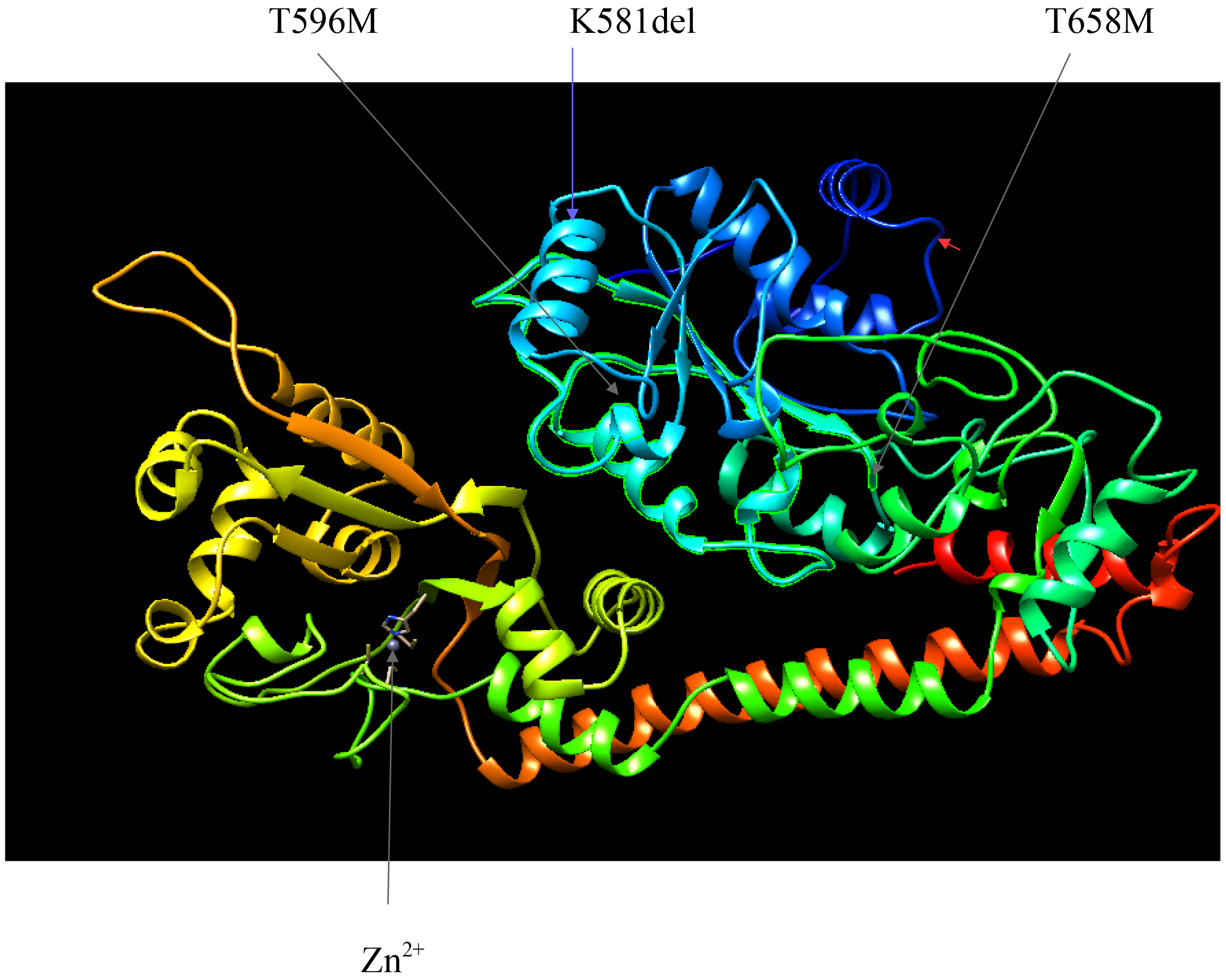
Screen-snipp of interactive model of *Cfam*-RECQL4 (ENSCAFG00000001569) folded onto the template crystal structure of human RECQL4 (Kaiser et al. 2017) PDB: 5LST. Presented ribbon model is rotated as in Fig. 1B in Kaiser et al. (2017). Zinc ion shown as blue sphere. The positions of residues altered by two missense varinats CTVT-T596M and CTVT-T658M are indicated by grey arrows (the DEAH box embedding helicase domain sub-region between CTVT-T596M and CTVT-T658M is higlighted in green). The position of inframe, splice-affecting CTVT-K581 deletion is indicated by blue arrow. Red arrowhead indicates the position of CTVT-R492 imposing the site entering frameshift onto predicted model. Image generated and colored with standard N-term. →C-term. code with UCSF Chimera ver. 1.12

## DISCUSSION

‘The clonal origins of tumour cell populations’, the concept developed by Nowell (1976), postulates the progression of most neoplasms results from an acquired genetic variants allowing for sequential selection of more aggressive sublines. The concept of clonal origins extended by Loeb (Loeb 1991; 2001; Loeb et al. 2003) implicates the survival or the life-span of the host is a rate limiting factor in considerations concerning the accumulation of the tumour genetic variants. The exceptional tropism of the transmissible tumours, however, permits for the almost limitless accumulation of the neoplastic variants. Consistent with the above notion, Murgia et al. (2006) established the modern-day CTVT does not appear to exhibit the severe mutator phenotype neither it does exhibit progressive chromosomal instability. Instead, Murgia and colleagues proposed the gross genetic and karyotypic rearrangements have occurred in transmissible Sticker sarcoma early in its emergence. To evaluate the above proposal, here we mined the catalogues of the patterns of genetic CTVT variants (Decker et al. 2015) concordant in two geographical transmissible tumour isolates reported by Murchison and colleagues (Murchison et al. 2014). We reasoned that the CTVT multiple-hit variant mining regime will uncover the out-of-phase candidates – tumour genes attaining the Knudsonian inheritance model - implicating the convergence of two (or more) somatic mutational events on a single gene. Here, we have described the relevant hetero-allelic candidate cases which arose with multiple-hit kinetics and provided the theoretical estimates of the likelihood of attaining the expected intragenic compound *trans*-heterozygous configuration.

### Identification of two tumour encoded RecQ-like DNA helicases accumulating the multitude of different mutational events

CTVT variant mining regime applied stringent criteria to identify Sticker sarcoma genes of oncologic relevance. While two critical identifications of canine RecQ-like DNA helicases are discussed below, few steps proceeding the acquisition of mutiple-hit out-of-phase candidates to final list demand a comment. **(*i*)** A number of canine multiple-hit out-of-phase candidates, identified at the extreme end of the spectrum of mutational load, are characterized by the gene-size bias. Amongst the most intensely mutated CTVT genes, identification of the putative canine titin ortholog provides an instructive example. Titin encoded by CTVT cells have accumulated a total of eight different mutations. *TTN*(enscafg00000014025) SNV polymorphisms listed with Decker set include three different inframe stop codon gaining events, two independent frameshift variants, splice acceptor variant in addition to two minor inframe variations (insertion and deletion respectively; details given in Table S2 by Decker et al. 2015). We wish to suggest that, assuming autosomal copy number equal (or greater) to 2n, in the absence of haplotype phasing information, the unambiguous and accurate estimate of the overall cumulative severity of given composite allele constellation pose a particular analytical challenge. For example, frameshift mutations might revert by the occurrence of a second complementary frameshift in *cis*- or other alterations involving e.g. splicing patterns of the same allele. This ‘haplotype phasing conundrum’ could be avoided if conventional sequencing of large recombinant clones of CTVT genomic DNA would be employed (particularly when the contamination with wild-type, host-derived, templates would be expected from nucleated stromal cells or leukocytes infiltrating the tumour - indeed estimated at ~15% of sequenced DNA in original report by Murchison and colleagues). We note the postulated analytical challenge may apply to other multiple-hit out-of-phase candidates. Clearly higher than 2n copy number is expected to contribute to the overall diversity of possible allelic distributions and increase the chance for retaining of the entirely intact wild-type copy. Moreover, stable polyploid CTVT forms are expected to further distract from the above expectations. Anyway, the patterns of recurrent mutations involving *TTN* were described elsewhere (e.g. Welch et al. 2012; Tan et al. 2015) in malignancies affecting humans and are consistent with background events affecting the size-biased genes (however an alternative interpretation have been suggested by Greenman et al. 2007). **(*ii*)** A considerable fraction of candidate cases which arose with multiple-hit kinetics represented ‘second-hit’ out-of-phase candidates and converged two SNVs on a given CTVT gene. Provided the theoretical estimates (see Theory and Results sections) as well as practical considerations those candidates were removed *a priori* with the stringently applied filtering criteria for selection of canine genes of oncologic relevance. This biased selection, likely containing gaps reflecting the limitations of the small animal oncology literature or shortcomings to our knowledge, was therefore based on canine-human comparative oncology (inferred from OMIM) and ultimately focused on genes causally implicated in established heritable cancer predisposing syndromes. We note this step might have not been entirely rational given a number of factors arbitrarily excluded from further SNV analysis. Some of those canine factors of oncologic relevance were affected by structural variants (SVs; listed separately in Decker set). Examples include canine tumour genes *ALK* (ENSCAFG00000005297) and *KIT* (ENSCAFG00000002065) affected by SV in addition to SNVs (see Table IV and Fig. 2 for other relevant examples excluded *a priori.*, but not affected by SV listed with Decker set e.g. *EGFR, RALB, RET*, and *RAD54B* [ENSCAFG00000009179]). The structural variants affecting tumour genes can contribute the equivalent of the second-hit, implicating additional out-of-phase candidate cases which arose with multiple-hit kinetics, and efficiently attain the compound *trans*-heterozygous Knudsonian inheritance model. The above considerations and limitations, however, does not change our overall conclusion concerning the postulated contribution of canine RecQ-like helicases to discussed below cellular phenotypes of transmissible Sticker sarcoma.

### Cellular phenotypes of CTVT-RECQL4 and CTVT-BLM – initial considerations

Previous analyses have specified several CTVT loci involved in genome maintenance and DNA-repair (Decker et al. 2015) however the possible contribution of predicted canine RecQ-like DNA helicases to cellular phenotypes of transmissible sarcoma were unnoticed. Analysis of candidate out-of-phase cases which perceivably arose with multiple-hit kinetics specifically indicated CTVT-RECQL4 among most intensely mutated genes encoded by Sticker sarcoma. Considering the perceived cumulative severity of somatic composite heterozygosity detected at CTVT-RECQL4 we note SNV polymorphism listed in Decker set specify that four out of six non-synonymous variants (frame-shift entering the reading frame at 492 residue, in-frame splice-affecting variant K581 and two missense substitutions T596M, T658M) predict to alter catalytic RECQL4 helicase domain and remaining two missense variants (A1116T and G1133R) alter the conserved C-terminal region (see Table S2 in Decker et al. 2015, for details; Larizza et al. 2010, for comparisons with human RTSII causing variants). Neither of the above CTVT variants nor any other canine *RECQL4* helicase variants have been reported in available literature to best of our comprehension. Provided the exceptional mutational load into CTVT-RECQL4, in the absence of haplotype phasing information, we postulate the demand for a dedicated experimental follow-up concerning the dissection and verification of perceived contribution of individual variants to specific neoplastic traits. We note the above argument applies to CTVT-BLM and possibly other mutations of multiple-hit kinetics. Further, isolate-specific mutations will expectedly increase the overall allelic complexity in the tentative composite heterozygotes. Again, in the absence of haplotype phasing information, the cumulative impact of those variants on multi-domain CTVT-BLM protein, presently may not be properly evaluated.

Since human cancer predisposing heritable progeroid syndromes are outside of the scope of this description and their characteristics and spectrum were covered elsewhere in the literature (e.g. Siitonen et al. 2009; Larizza et al. 2010; reviewed in Bachrati & Hickson 2003) we focus the discussion on the factors of relevance in canine comparative oncology. Both human BLM and RECQL4 are regarded and classified as ‘chromosome breakage syndromes’ however the cases of combined deficiency have not been described. Cellular phenotypes of RecQ-like helicase deficiency covered in the cited literature include the chromosomal instability with various cytogenetic abnormalities. Cell lines established from Bloom syndrome (BS) patients were described as hyper-recombinogenic with gaps, breaks and structurally rearranged chromosomes including autosomal and Y chromosome loss (e.g. Shabtai & Halbrecht 1980, cited and discussed in Chester et al. 2006 with other relevant references). Cell lines established from RTSII patients represent variegated mosaic aneuploidies (Der Kaloustian et al. 1990) with isochromosome formation (Larizza et al. 2010). The characteristic feature distinguishing between chromosomal instabilities affecting BLM and RECQL4 deficient cells is the rate of sister chromatid exchange (SCE), increased by many-fold in BS but not RTSII. The experimental combined BLM/RECQL4 withdrawal reported (Sight et al. 2011), suggest that in the SCE prone - BLM (-/-) background, RECQL4 depletion contributes to the measurable increase in SCE formation (Fig.9 in Sight et al. 2012). The SCE assay however, has not been reported in CTVT neither elsewhere in the literature concerning the canine oncology, alas.

Considering the cellular phenotypes of RecQ-like DNA helicases deficiency, the sensitivity to treatments with DNA-damaging agents appear of possible relevance to CTVT. In particular chronic photosensitive component described in BS (recapitulated in BLM deficient cells) is reflected by the elevated SCE rates in BS cells, further increased in the response to ultraviolet-C exposure (Ababou et al. 2002, with relevant references). It has been proposed that UV sensitivity observed in BLM deficient BS derived cells could be attributed to photoadducts repair-deficiency and possibly some other impaired DNA repair mechanisms (Bachrati & Hickson 2003). While, UV sensitivity in RTS derived cells appears less well established (Shinya et al. 1993; Vasseur et al. 1999; reviewed in Larizza et al. 2010), RECQL4 has been implicated in a repair of UV induced damage (Fan & Luo 2008). The influence of combined RecQ withdrawal of BLM/RECQL4 on UV sensitivity, however, have not been carefully characterized (Singh et al. 2012).

Previous reports (Murchison et al. 2014 and Decker et al. 2015) specified the burden of CTVT mutational signatures consistent with the exposure of tumour cells to UV light-induced DNA-damaging photoadducts. Hence, we propose the RecQ helicase dependent repair-deficiency could have predilected Sticker sarcoma cells to UV induced photoadducts, and suggest thereof a preponderance of mutational signatures associated with UV exposure could be explained by the dominantly inherited susceptibility to UV light.

### Aneuploidy

Among the cellular neoplastic phenotypes of CTVT cells, possibly affected by multiply-mutated RecQ-like helicases, are traits perceivably contributing to generalized aneuploidy of Sticker sarcoma. Generalized aneuploidy perceivably may be influenced by mutant CTVT-RecQ-like helicase. The neoplastic traits associated with aneuploidy are widely recognized and covered in comparative oncology literature (e.g. Bloomfield & Duesberg 2016; Taylor et al., 2018, reviewed in Duesberg and Rasnick 2000; Rajagopalan & Lengauer 2004; Pathak & Multani 2006). While aneuploidy have been recognized in non-transmissible canine neoplasms, including mammary tumours (Rutteman et al. 1988, Hellmen et al. 1993; Cassali et al. 2007), hemangiopericytoma (Kang et al. 2006), melanoma (Bolon et al. 1990), mast cell tumours (Ayl et al. 1992) and osteosarcoma (Fox et al. 1990; Maeda et al. 2012), as a parameter of clinical relevance (Lucroy 2008) that may predict aggressive tumour behaviour – however, the specific connections to mutant RecQ-like helicases are still to be established (Fan & Khanna 2015). Collectively, the proposal concerning the perceived contribution of the multiple-hit RecQ-like candidates to CTVT cellular phenotypes, is thereof originally specified by our theoretical analysis of variants listed in Decker set. The above proposal concerning the involvement of the multiply-mutated RecQ-like helicases in naturally occurring canine transmissible tumour, is further supported by the independent description of the experimental work involving the engineered mouse models. Considering the phenotypes of canine RecQ helicase deficiencies in mammals other than man (where RecQ helicases were causally implicated in cancer predisposing heritable syndromes; particularly well studied Bloom syndrome and not as well documented type II Rothmund-Thomson syndrome; described and reviewed elsewhere in cited literature and not specifically discussed here) mouse models engineered with embryonic stem cell technology appear of particular relevance. Mann et al. (2005) description of a viable homozygous mouse model of murine *Recql4* (-/-) variant provided an independent evidence for coupled aneuploidy and cancer predisposition. Cancer predisposition in a viable homozygous mouse model of murine *Blm*(-/-) was described by Luo et al. (2000) and engineered *Blm* conditional alleles were described by Chester et al. (2006). Remarkably the later description of conditional *Blm* allele established the engineered RecQ deficient neoplastic traits are biased towards aneuploid tumour karyotype and secondary genomic instability. While combined viable doubly homozygous *Blm* (-/-); *Recql4* (-/-) murine lines have not been described (neither the case of *Blm* / *Recql4* second-site non-allelic non-complementation), we suggest such combined RecQ deficiency would model the combined CTVT-RECQL4 / CTVT-BLM deficiency most accurately. Importantly aforementioned combined *RECQL4* / *BLM* withdrawal was reported by Singh et al. (2012) in cell lines, to conclude both physical interaction and functional cooperation, between human RecQ helicases RECQL4 and BLM, in broadly defined maintenance of the genome stability. We suggest the proposed cooperative actions of CTVT RecQ-like DNA helicases could be relevant interpreting whether contributing RecQ dependent karyotypic traits, respond to selective pressures that preserve the aneuploid genomic structure of transmissible Sticker sarcoma.

Considering further the possible contribution of multiply-mutated RecQ helicases to traits maintaining the neoplastic aneuploidy of CTVT genome, we proposed one of the three unique CTVT-BLM variants: G973E, constitute the catalytic-null. Catalytically impaired CTVT-BLM variant may result in BLM haploinsufficiency. *Blm* haploinsufficiency was suggested in a mouse model: Goss et al. (2002) described enhanced tumour formation in viable *Blm* (+/-) heterozygous mice, an observation consistent with the notion of Gruber et al. (2002) reporting the two-fold increase in the occurrence of colorectal cancer in human *BLM^Ash^* carriers. Those studies concluded that only one mutant allele of *Blm/BML* could be regarded as a modifier of cancer predisposition, hence a relevant factor in the maintenance of neoplastic aneuploidy. Provided G973E is concordant in two sampled CTVT isolates, at least one allele of CTVT-*BLM* could be regarded as inactivated in a prototype tumour regardless of the overall heteroallele configuration. While aware of difficulties in the evaluation of cumulative pathogenicity of variants that converged on CTVT-*BLM*, we remark, the above argument is further strengthened by notion of others (Neff et al. 1999; Wang et al. 2001; Yankiwski et al. 2001) proposing that certain allelic variants of *BLM*, including those regarded as catalytic-nulls, could possibly contribute to dominant traits. In particular certain allelic variants of BLM, were demonstrated to miss-localize the wild-type BLM perceivably by incorporation of catalytically inactive variants into homo- and hetero-meric protein complexes engaged by BLM. *BLM* alleles contributing to dominant traits were proposed to further sequester the function of BLM dependent DNA-repair complexes and thereof affect the genomic stability. We suggest, that the above scenario could apply to multiply-mutated CTVT-*BLM* and possibly explain how heteroallelic variants encoded by transmissible sarcoma genome might contribute to the maintenance of the aneuploid CTVT karyotype.

Clearly other than canine RecQ-like factors encoded by CTVT might contribute to aneuploid karyotype. Decker et al. (2015) specified somatic mutations in DNA repair and genome stability genes as likely contributing to CTVT genomic disarray. We emphasize that certain factors other than RecQ-like helicases, specified in the description by Decker and colleagues, including *ATM, BRCA1, FANCD1/BRCA2*, and *MRE11* were enriched with mining scheme applied here i.e. converged two (or more) mutations on any given gene and thereof perceivably arose with multiple-hit kinetics. Those candidate out-of-phase multiple-hit identifications, are of possible relevance as, encode for factors known to interact with BLM either directly or indirectly e.g. BRCA1 and MRE11 (Wang et al. 2000), FANCD1/BRCA2 via multiprotein nuclear complex with FANC proteins (Meetei et al. 2003), and / or functionally e.g. subjected to post-translational phosphorylation by ATM (Beamish et al. 2002, reviewed in Bachrati & Hickson 2003, Cunniff et al. 2017). We note here that multiply mutated CTVT-ATM and CTVT-BRCA1 represent second-hit out-of-phase candidates accumulating only two seemingly mild missense substitution variants. Those appear to group with multiple-hit candidate CTVT-FANCD1/BRCA2 (two missense variants and inframe deletion affecting single codon) as variants of unknown significance demanding further verification as possibly functionally synonymous with wild-type. Our analysis implicated yet another previously unnoticed DNA-repair factor CTVT-FANCD2 centrally involved in the FANC pathway and genome stability maintenance. In contrast to CTVT-FANCD1/BRCA2 mentioned by Decker et al. (2015) and affected by relatively mild polymorphisms (above), CTVT-FANCD2 accumulated three missense variants (V615I, D751G and S1421F) in addition to inframe stop codon gaining SNV (E1303*) consistent with the premature translation termination. Provided known association of BLM with FANC complex (Meetei et al. 2003, Payne & Hickson 2009) the later report by Naim & Rosselli (2009) specified the functional associations of BLM and FANCD2 in preventing the micro-nucleations and reducing the chromosomal abnormalities under the replicative stress. The proposed cooperative role of BLM and FANCD2 in preventing aneuploidy seems relevant given both *BLM* and *FANCD2* encoded by aneuploid genome of transmissible sarcoma accumulated multitude of different mutations.

Two other factors involved in DNA-repair and genome maintenance described by Decker and colleagues (2015) include CTVT-MLH1 and CTVT-TP53. Based on the model drawn from interactions mapped in human cells, MLH1 and TP53 are known as functional partners directly interacting with BLM (Wang et al. 2001; Garkavstev et al. 2001; reviewed in Bachrati & Hickson 2003). While neither CTVT-MLH1 nor CTVT-TP53 were enriched with our multiple-hit mining scheme the later identification is of possible relevance, provided postulated interaction and cooperation between TP53 and RecQ helicase BLM. Considering the factors involved in maintenance of the aneuploid genomic structure of CTVT, based on the models proposed to operate in neoplasms in systems other than canine, Fujiwara et al. (2005) demonstrated mutant *p53* (-/-) promotes the formation of aneuploid malignancies via mechanisms involving the polyploid intermediates. Further, Vitale et al. (2010) described the generation of aneuploid cells descended from multipolar asymmetric divisions of tetraploid precursors under the absence of *p53* (reviewed in Margolis 2005; Castedo et al. 2010). Given the demonstration of CTVT forms regarded as ‘stable polyploids’ (Thorburn et al. 1968 and Sellyei et al. 1970) and provided the monoclonal origins of transmissible Sticker sarcoma (Murgia et al. 2006), the above associations are of possible relevance in considerations pertaining to the perceived transitions between aneuploid and polyploid variants. In the absence of direct evidence, we note others (reviewed in Rajagopalan & Lengauer 2004; Krajcovic & Overholtzer 2012) specified mechanisms leading to neoplastic aneuploidy involving polyploid intermediates. Whether above mechanisms involved in genome-dosage alterations account for the cellular phenotypes of CTVT cells and if those do relate to DNA-repair/genome maintenance factors including CTVT-TP53 and canine RecQ-like DNA helicases, remains to be demonstrated (Stockmann et al. 2011). We further note CTVT-TP53 is affected by the structural variant of unusual complexity - the local intra-chromosomal inversion on canine chr 5 - creating the reciprocal fusions (involving the CTVT-TP53 split into two, respectively N- and C-terminal, fragments reciprocally joined through mid-intronic segments) with parts of another CTVT gene SHBG (Decker set). On this occasion we also note, in addition to the typical aneuploid transmissible venereal tumours involving ‘organs concerned with the reproduction’, clinically manifesting Sticker sarcomas include atypical extra-genital presentations e.g. cutaneous (Ferreira et al. 2017), mucosal of oral and nasal cavity (Razaei et al. 2016) as well as disseminated (Park et al. 2006) and otherwise generalized (Shafiuzama et al. 2015) forms. Sticker sarcoma is regarded clinically as conditionally malignant neoplasm with the occasional emergence of metastatic lesions involving lymph nodes and internal organs (reviewed in Das & Das 2000). Until now the above atypical presentations have not been systematically assessed (see footnote in method section) either evaluated to modern genetic standards, precluding additional variant analyses in aneuploid and stable polyploid CTVT forms. Together, we suggest investigations of mutational patterns and underlying aneuploid genomic architectures of altered tropism isolates, malignant and metastatic forms etc., will inevitably permit for critical assessment of the involvement of RecQ-like helicases and possibly other multiply-mutated factors.

### Loss-of-heterozygosity and mitotic recombination

Neoplastic aneuploidy is often accompanied by two related phenomena: loss-of-heterozygosity and mitotic recombinations. Loss-of-heterozygosity (LOH) appears of broad relevance, owing to the fact it may efficiently contribute an equivalent of the second-hit (Cavenee et al. 1983) and expectedly alter the allelic configurations of a tumour genes. Extending on the earlier statement (Murchison et al. 2014), concerning a large proportion of aneuploid CTVT genome contracting LOH events, report of Decker et al. (2015) provided genome-wide map of regions of the Sticker sarcoma genome affected by sequential LOH waves. Our results i.e. analysis implicating multiply-mutated CTVT RecQ-like helicases appears of relevance - provided the demonstration by LaRocque et al. (2011) that BLM withdrawal leads to a substantial increase in the efficiency of LOH, presumably resulting from somatic crossing over between homologs. Results of LaRocque and colleagues are consistent with increased frequency of spontaneous LOH reported in mouse (Yusa et al. 2004, Guo 2004) and human (Langlois et al. 1989; German 1993; Ellis et al. 1995a) BLM-deficient cells. Furthermore, mechanisms inferred from murine (Luo et al. 2000) and fish (Xie et al. 2007) models suggest that certain RecQ-like helicases may act as tumorigenesis suppressors by preventing LOH through the suppression of ectopic recombination between homologous chromosomes. Additionally, somatic intragenic recombination events altering the *trans-cis* allelic configuration of compound *BLM* heterozygotes (resulting in the restoration of a wild-type copy via *trans-cis* allelic transition) were described in cells isolated from Bloom syndrome patients (Ellis et al. 1995b; Ellis et al. 2001; German 2004; reviewed Weksberg 1995 and elsewhere in cited literature). Collectively, the widespread frequency of homozygous variants, indicating altered allelic segregation in CTVT genome, could be taken as indirect evidence for increased somatic recombination in Sticker sarcoma cells. The segmental and sequential nature of the widespread LOH described in CTVT by Decker and colleagues argues in favour of this proposal. While the alternative, albeit mutually non-exclusive, explanation for the relatively low prevalence of heterozygous variants in founder CTVT postulated by Murchison and colleagues might exist (e.g. implicating the uni-parentally disomic CTVT origins with possible involvement of genome-wide chromosomal non-disjunction; plausible in particular case when the prototype CTVT would have erupted from a neoplastic form of unusual conceptus - such us gestational trophoblastic malignancy) we note that the later explanation remain speculative in nature. Given the broadly defined role of RecQ-like helicases acting as illegitimate recombination suppressors, the involvement of multiply-mutated CTVT-RecQ-like helicases, i.e. *BLM* and other cooperating multiple-hit candidates, together might provide the mechanistic explanation concerning the factors contributing to the maintenance of generalized aneuploidy of CTVT genome.

Considering the postulated CTVT *RecQ*-deficiency model perceivably implicating the mitotic recombination, in the evolutionary perspective: we emphasise, the somatic crossing-over between homologous chromosomes may efficiently bypass seemingly inexorable effects of the Muller’s Ratchet (Muller 1964; Felsenstein 1974). In that perspective postulated CTVT *RecQ*-deficiency (inferred from theoretical analysis of multiply-mutated CTVT-*BLM* and CTVT-*RECQL4* and other *RecQ* auxiliary factors) might represent an adaptive neoplastic trait, extending on the proposal of Olson (1999) - ‘when less is more’ - concerning the evolutionary role of gene loss.

## Supporting information

Supplemental PDB file 1

Supplemental PDB file 2

Supplemental PDB file 3

Table files III-VI

Table files VIII

**Table III.** Cases of most intensly mutated CTVT genes. Table supplied as an Excel file.

**Table IV.** CTVT doubly-mutated genes of oncological relevance. Table supplied as an Excel file.

**Table V.** CTVT multiply-mutated genes of oncological relevance. Table supplied as an Excel file.

**Table VI.** Human orthologs of muliply-mutated candidate CTVT genes and associated hereditary neoplastic syndromes in man. Table supplied as an Excel file.

**Table VIII.** CTVT-RecQ helicases - mapping of structural variants. Table supplied as an Excel file.

## Footnotes

1 )Original description provides the relevant photographs (Fig 1A in the description by Murchison and colleagues) documenting the location of sequenced tumours with respect to body parts of tumour affected hosts: picture of tumour 24T localized in caudal area clearly involving vulva (and presumably internal part of female reproductive organs) in 24H host and anterior presentation for the 79T tumour affecting male host 79H (presumably an atypical, extra-genital tumour presentation).

## Abbreviations

CTVT: Canine transmissible venereal tumors
BS: Bloom syndrome
RTS, RecQ: Rothmund-Thomson syndrome
SNV: Single-nucleotide variant
CNV: Copy number variant
SV: Structural variant
LOH: Loss of heterozygosity
SCE: sister chromatid exchange
UV: ultraviolet
Kb: kilobase
Mb: megabase

